# On the characterization and interpretation of phenotypic ellipse geometry, with examples from *Canis* and sigmodontine rodents

**DOI:** 10.64898/2026.07.21.739591

**Authors:** F. Robin O’Keefe

## Abstract

Data on the shape of a group of organisms can be conceptualized as forming a point cloud in the multivariate space of measurement. This is literally true for traditional linear measures, while in a geometric morphometric context the cloud resides in Kendall’s shape space, tangent to the true shape space. Regardless of the method of construction, the topology of this point cloud, or phenotypic (hyper)ellipse, is a beguiling target for evolutionary analysis. Reordination of the axes will not change the geometry of this phenotypic ellipse, and the notion that its geometry carries a meaningful biological signal is an old idea; but the character of this signal is often elusive. This paper explores the application of the most commonly used parameter designed to summarize differences in phenotypic ellipse geometry (relative eigenvalue variance, or V_rel_), and demonstrates that it is incapable of differentiating between several plausible ways in which phenotypic ellipse geometry might differ among species, because it confounds three separate parameters necessary to describe the ellipse. Two example data sets are analyzed to illustrate variability in phenotypic ellipse geometry and draw conclusions about observed differences. The first case compares wolves to domestic dogs, and replicates previous findings of much greater variance yet tighter integration in dogs. This calls into question the simple model of a single peak in the fitness landscape of dogs. The second example comprises geometric morphometric landmarks from the jaws of a clade of sigmodontine rodents, and allows comparison of ellipse geometry in a phylogenetically controlled setting with qualitative ecological categories. Three parameters are found to vary in concert along a grade of most to least ecologically specialized: the phenotypic variance, the effective rank (dimensionality), and the degree of covariance (V_rel_ and related metrics). Use of all three of these quantities to characterize the geometry of the phenotypic ellipse is advocated, as all are necessary to characterize how variance is distributed in the ellipse in different taxa. The phenotypic ellipse geometries illustrated here appear to reflect something of the geometry of the adaptive peak upon which each taxon sits; or that they represent (aspects of) the mapping function of adaptive peak to phenotypic ellipse, in ways first predicted by Simpson in the twentieth century.

---

> *“Serious studies of Adaptive Landscapes need to integrate historical records (via… cladistics), functional ecology, as well as morphology—a splendid area for interdisciplinary work, much of which remains to be done.”* --Pigliucci, 2012 p. 36.

## Introduction

The goal of this paper is to investigate the notion that the geometry of the space occupied by a species’ phenotype reflects something of the adaptive landscape upon which it rests. As stated above by Pigliucci (above, 2012), this question sits at the intersection of several evolutionary lines of thought, and so is inherently challenging, as three distinct types of data are required: a measure of phenotype, a measure of ecological function or niche breadth relevant to the phenotype, and control of phylogenetic autocorrelation. This paper is concerned primarily with one of these data types, namely the geometry of the space occupied by a species’ phenotype. A mathematical framework is developed for the full characterization of the topology of this space, here termed the *phenotypic (hyper)ellipse*. This framework treats explicitly three of the four parameters necessary to characterize the topology of the phenotypic ellipse: its volume; its shape; its dimensionality; and its orientation. The framework is then applied to two illustrative data sets to demonstrate its utility, with elementary but accurate treatments of ecological breadth and phylogenetic autocorrelation. The results presented below demonstrate that phenotypic ellipse geometry varies significantly among closely related taxa, and further that the observed differences have plausible (and testable) biological interpretations. It is further argued that measures of “overall modularlity and integration” (Goswami and Polly, 2010; Pavlicev et al., 2009a; O’Keefe et al., 2022) confound several distinct attributes of the phenotypic ellipse, and that this may account for the known difficulty in their biological interpretation (Goswami and Polly, 2010; see discussion in Watanabe, 2022). This paper concerns many of the of the topics and techniques developed for the analysis of phenotypic modularity and integration, but the goal here differs. Modern research on integration is concerned with the assembly and disassembly of patterns of covariance within the phenotypic ellipse, (Klinginberg, 2008), while this paper is concerned with the external topology of the ellipse.

### The Phenotypic Ellipse

The characterization of phenotype is a multivariate problem. Biological shapes are complex, and measurement strategies designed to capture this complexity involve multiple variables, be they ‘traditional’ linear measures (Marcus, 1990) or landmark coordinates (Zelditch et al., 2012). Principal components analysis (PCA) is standard practice for reducing the dimensionality of the resulting covariance matrix, where reordination is used to yield a few composite variables embodying most of the variance in the original matrix (Goswami and Polly, 2010). These are then used to summarize the data for subsequent analysis (Goswami et al., 2019; for an introduction see Reyment, 1991). In most biological studies it is the scores on the first few principal components that are of primary interest, as they embody of the major axes of shape variation in the sample (Drake and Klingenberg 2010; Goswami and Polly, 2010; O’Keefe et al., 2016; Evin et al., 2025). PCA yields a new covariance matrix of orthogonal composite variables (eigenvectors) with zero covariation on the off-diagonals and a vector of associated variances (the eigenvalues) on the diagonal. The first eigenvalue is generally the largest, the second is smaller, and subsequent eigenvalues decrease rapidly in magnitude (Marcus, 1990). The characterization of the phenotypic ellipse developed here rests explicitly on PCA as applied to geometric morphometric data, namely landmark coordinate data after Procrustes superimposition (Bookstein, 1991).

Given a sample of *n* individuals measured over *p* phenotypic traits, we may envision the individuals as occupying a cloud, or hyperellipse, in the space defined by the traits (Van Valen 1974; Watanabe, 2022). At an abstract level, this cloud may be characterized uniquely by four parameters: 1) the orientation of the principal axes of the cloud relative to the trait axes; 2) the volume of the cloud; 3) the dimensionality of the cloud, (usually less than the full rank implied by the measurement scheme; O’Keefe et al., 2022); and 4) the degree of correlation among traits. A set of highly correlated measures will yield and longer and skinnier cloud than a set of less correlated measures, whose cloud will be more oblate (Watanabe, 2022).

Conceptualizing the phenotypic ellipse in this way makes a number of assumptions. The traits are assumed to be continuous and real (i.e. not categorical); each trait is further assumed to be normally distributed; and the correlations between traits are assumed to be adequately represented by linear models. Granted these conditions, we may then ask what, if anything, the topology of this ellipse can tell us biologically. Barriers to interpretation are legion. As a rule, phenotypic trait spaces are tailored closely to the organisms under study, so that trait spaces are highly specific to their taxa, and it is simply not possible to plot (for example) an insect wing into a phenotypic space designed for bird wings. This limitation can be overcome by narrowing the focus of organisms studied and/or using a more general trait space, but comparative morphometric studies must always strike a balance between taxic breadth and anatomical detail. Yet given appropriate selection of taxa and measurement scheme, comparison is possible, and in fact the specific angles that principal axes make with each other, and with the original trait axes, can be calculated explicitly (O’Keefe et al., 1999). These angles are embodied in the coefficients of the eigenvectors produced by PCA. While interesting, the specific orientations of phenotypic ellipses are not treated specifically in this paper.

Of the four parameters listed above, volume is the easiest to measure, as it is simply the sum of the constituent trait variances (termed the phenotypic variance by Drake and Klingenberg, 2010), or equivalently the sum of all eigenvalues after PCA. The classic study on wild and domestic canids by Drake and Kingenberg (2010) explores the phenotypic contrast between gray wolves and their direct descendants, the domestic dog. Dogs are found to possess about seven times as much phenotypic variance as wolves, and this drops only to five times when the greater size variance of dogs is regressed out. The volume of the phenotypic ellipse is therefore much greater in domestic dogs than in the wolves they evolved from, and we might posit that an ecological release of stabilizing selection in dogs to account for this. However this hypothesis does not bear further examination; another main objective of that paper was to compare the degree of integration between the neurocranial and viscerocranial modules in dogs vs. wolves. Wolves were found to exhibit modest modularity, while dogs are much more tightly integrated (Drake and Klingenberg, 2010). That paper attributes the tighter integration in dogs to their much greater variance; however, this conclusion follows only if the additional variance is confined to the largest eigenvalues, and rank does not increase. As described below, an overall increase in variance should result in an *increase* in modularity if the covariance structure does not change, the opposite of what is observed. One goal of this paper will be to revisit this study using the recent data set of Evin et al. 2025, in an effort to shed light on this puzzling observation.

### Ellipse Shape: Eigenvalue Dispersion and the Relative Eigenvalue Variance

> *Over several generations of evolutionary and developmental biologists, ever since Olson and Miller’s pioneering work of the 1950’s, the concept of “morphological integration” as applied to Gaussian representations N(μ, Σ) of morphometric data has been a focus equally of methodological innovation and methodological perplexity.* – Bookstein, 2022; p. 342.
>
> *However, the use of these indices [eigenvalue dispersion metrics] has been criticized for a lack of clear statistical justifications; it has not been known—or not widely appreciated by biologists—exactly what they are designed to measure, beyond the intuitive allusion to eccentricity… –* Watanabe, 2022; p. 5-6.

As discussed at length by both Bookstein (2022) and Watanabe (2022), the use of summary indices to characterize the eigenvalue distribution has a long and imposing history, a history that should (and does) inspire trepidation in the mind of any researcher wishing to contribute meaningfully to the edifice. Yet work remains to be done; the two papers cited above both treat eigenvalue dispersion indices exhaustively but reach diametrically opposed conclusions as to their utility. Watanabe (2022) offers a broad and comprehensive theoretical footing for the most common of these indices, the relative eigenvalue variance (or V_rel_, Table 1; Pavlicev et al., 2009a; Goswami and Polly, 2010; for a thorough theoretical introduction see Najarzadeh, 2019). The V_rel_ parameter is meant to quantify the premise that matrices with stronger covariation will have greater eigenvalue variance, while those with less covariation will have less variance. Watanabe (2022) frames V_rel_ on an intuitively appealing test of sphericity of the phenotypic (hyper)ellipse, and utilizes a simulation study to establish its statistical behavior. Watanabe’s equivalence of the V_rel_ to a test of sphericity is a useful starting point for visualizing the geometry of the data cloud whose properties V_rel_ is designed to measure (Watanabe, 2022). Given a matrix of *n* samples measured over *p* variables of equal variance, the special case of zero correlation between all variables will yield a *p*-ball, or a hypersphere of points of spatial dimension *p* (this is not the full shape space but Kendall’s shape subspace, which is tangent to the full shape space; Zelditch et al., 2012; Bookstein, 1991). As covariance among variables increases, the ball will squeeze more tightly into a hyperellipse whose aspect ratio is proportional to the strength of covariance (Van Valen, 1974; Cheverud et al., 1983; Conaway and Adams, 2022). A covariance matrix derived from such a hyperellipse will retain the variances on the diagonal, while the off-diagonal covariance terms become non-zero. This is the norm, as biological systems are in general highly correlated (Cheverud, 1982; Pavlicev et al. 2009a; Watanabe, 2022; Bookstein, 2022), and it is the strength of this covariation, and hence the skinniness (Or ‘tightness’, to use Van Valen’s term) of the hyperellipse that V_rel_ and related indices are designed to measure. But it is not immediately clear what this tightness means biologically, beyond Watanabe’s “intuitive allusion to eccentricity.”

**Table 1.**
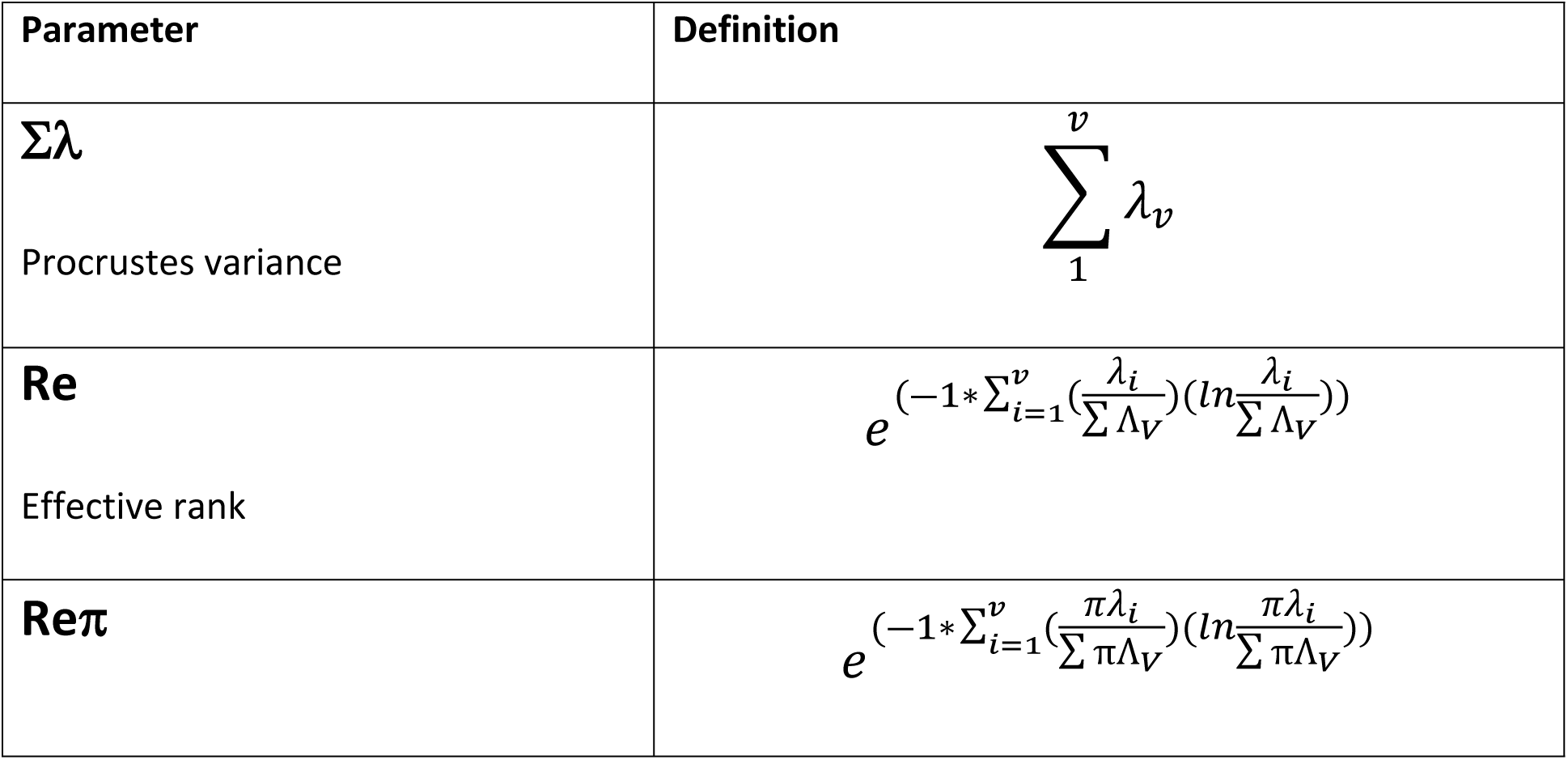
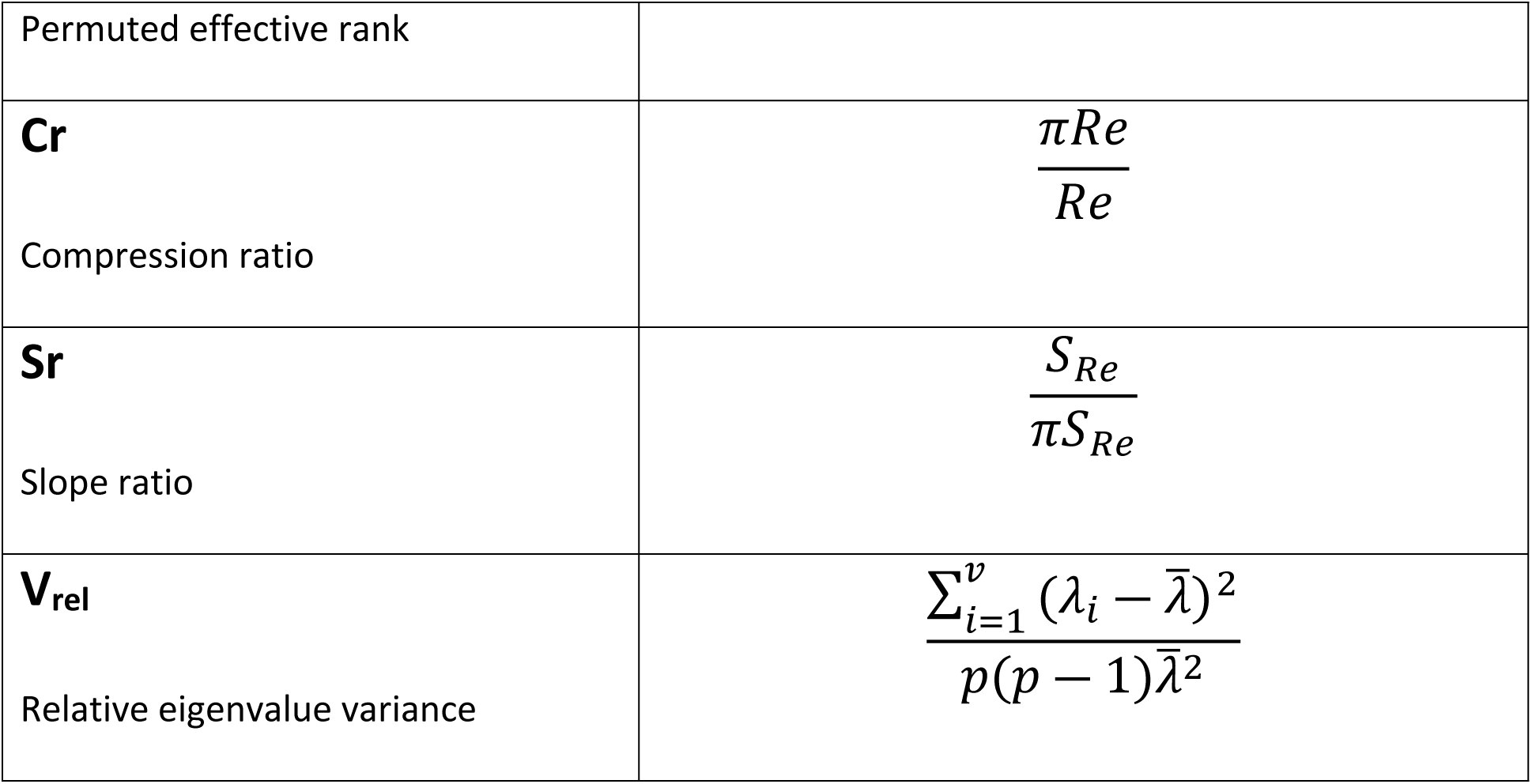
Quantities defined and applied in this study. All expressions are defined in the Introduction.

By contrast, Bookstein’s 2022 paper reads as an indictment of V_rel_ and related indices. His main thesis is that the features of biological interest in covariance matrices are internal patterns of covariance, and that these patterns are lost when a summary index is calculated. Bookstein makes several trenchant methodological points, the first being that V_rel_ is a function where the mean eigenvalue appears in the numerator, and hence information concerning the magnitude of phenotypic variance is lost. His second point is that the use of the correlation matrix (**R**) rather than the covariance matrix (**C**) is unwise, because the *Z* transform used to create the correlation matrix obscures (biologically real) differences in the magnitudes of the input variables (for further discussion of the use of **R** see O’Keefe et al., 2002). Both points are easily addressed; the phenotypic variance is easy to calculate, and one may restrict calculation of V_rel_ to **C** (defined as a covariance matrix derived from a matrix of geometric morphometric data after Procrustes superimposition). But Bookstein’s core challenge remains: do eigenvalue dispersion indices in general, and V_rel_ in particular, express anything meaningful biologically? He believes not, and as quoted above Watanabe concurs that biological interpretation is, at least, elusive. This paper attempts to throw some light on these questions through the lens of information theory and associated permutation tests, applied to a well-studied model system in sigmodontine rodents. The conclusion of this exploration is that a straightforward biological interpretation of V_rel_ is not possible, because it confounds several distinct phenomena needed to characterize fully the geometry of the phenotypic ellipse. Further exploratory analysis with both rats and dogs demonstrates that full knowledge of the topology of the phenotypic ellipse allows for hypotheses concerning the conformation between the phenotypic ellipse and the fitness surface it rests upon.

### Dimensionality: Matrix Rank

Van Valen (1974) noted that phenotypic hyperellipses may vary in their dimensionality as well as their relative degree of covariance, and suggested a way to measure this, although it has not seen wide use. However, awareness has grown that phenotypic matrices in general, and those derived from geometric morphometrics in particular, are usually rank deficient (O’Keefe et al., 2022). A parameter based on the eigenvalue distribution must therefore cut off a tail of trivially small eigenvalues to be useful. Pavlicev et al. (2009b) suggest a bootstrap procedure to accomplish this, while the current version of the R package **geomorph** (Baken et al., 2021; Adams et al., 2025) takes only eigenvalues “greater than zero” to make this cutoff (Conaway and Adams, 2022; the procedure for determining equality to zero is not specified in the documentation). The method used by O’Keefe et al. (2022) to characterize the effective dimensionality, or *effective rank*, utilizes the spectral entropy (Shannon, 1948) of the eigenvalue distribution to quantify the degree to which the nominal rank of such matrices is greater than the effective rank. the Shannon entropy *E_s_* of a vector of eigenvalues **Λ** derived from PCA of **C** is given as:

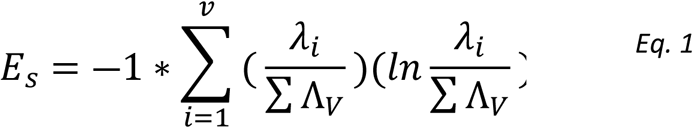

The effective rank is then calculated by raising *e* to the power of the Shannon entropy:

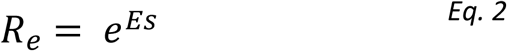

Effective rank calculated in this way allows one to establish one good estimate of the significant number of principal axes, or equivalently the dimensionality, of a covariance matrix. An excellent introduction to this issue, relevant literature, and R code involved in identifying rank deficiency in biological, ecological, and neuroscientific covariance matrices may be found in Del Giudice, 2021.

The nominal, full rank of **C** (= *v*) is vulnerable to loss in matrix rank from at least three different sources: 1) a lack of adequate sample size (n); 2) loss of degrees of freedom due to Procrustes superimposition of landmark coordinates; and 3) biological covariance (O’Keefe et al., 2022). Only the third of these is of biological interest. Procrustes superimposition of landmarks sacrifices 4 (2D) or 7 (3D) degrees of freedom (Zelditch et al., 2012), rendering the last 4 or 7 eigenvalues of a PCA of Procrustes shape coordinates formally equal to zero. Additionally, the required sample size for phenotypic measures is suggested to be least 10 times the number of variables (Grabowski and Porto, 2017), meaning that data with fewer samples than 10*v* will be rank deficient. This drop in matrix rank is large and must be considered; for instance, two samples of La Brea carnivore jaws (both comprising 14, 2D landmarks) have a nominal rank of 24, but observed effective ranks of only 11.36 and 12.32, respectively (O’Keefe et al., 2022). If half of the eigenvalues in a distribution are meaningless, then the biological signal in any moment of that distribution will be obscured. This problem has been shown empirically, with Conaway and Adams (2022) demonstrating that none of the proposed metrics of phenotypic integration they tested performed well, with the sole exception of V_rel_. The few integration metrics that consider both dimensionality and strength of covariance (e.g. Van Valen, 1974, and O’Keefe et al., 2022) attempt to combine the two phenomena into a single metric; however, Conaway and Adams (2022) find O’Keefe et al.’s suggested index to be ineffective. While it is provocative that *Aenocyon* and *Smilodo*n differ significantly in rank when both nominal rank and sample size are held constant, attributing any biological meaning to this difference is hobbled in that study by (slight) differences in landmark configuration and (great) differences the amount of time averaging between taxa. To explore possible variability in both covariance and dimensionality at the same time, a more controlled species comparison with identical landmark schemes, sample sizes, ecological characterizations, and a phylogenetic context is needed (Pigliucci, 2012). A major objective of this paper is analysis of such a case from sigmodontine rodents; this case demonstrates clearly that both covariance and dimensionality vary significantly between closely related species.

### Expectations for Ellipse Geometry

Because phenotype is the interface between genotype and the environment (Tamagnini et al., 2021), we may posit that the hyperellipse reflects something of the underlying geometry of the fitness landscape upon which it resides (Wright, 1932; Dietrich and Skipper, 2012; expressed as conformation between environmental (*E*) and phenotypic (*P*) sets by Cheverud, 1982). This is an old idea, reaching back to the mid-20^th^ century evolutionary synthesis and presented elegantly, if qualitatively, by Simpson (1953). Given a fitness surface with a single, static peak that coincides with an ideal shape, Simpson expects that a population under more intense selective pressure around this peak will display less variance than one under less intense selection (Figure 1). This is a testable prediction: that species with more specialized ecological preferences will vary less than those with broader ones, and this is often observed (for instance Van Valen, 1965). This hypothesis is termed the Specialist-Generalist Variation Hypothesis (SGVH) by Li et al., 2014. Those authors describe a case of this effect in the genetics of *Caenorhabditis*, and their survey of a diverse array of other organisms suggests the SGVH holds broadly. A direct test of varying selection regimes on wing phenotype in successive *Drosophila* generations demonstrated that populations under more intense selection show decreased variation (Pélabon et al., 2010). Obviously the caveats are many; the literature on the challenges of ecological niche delineation is vast, and phylogenetic history is at least as important as environment in molding shape variation (Maestri et al., 2018). Further, the presence of shape covariance (and genetic pleiotropy) can both constrain and canalize the directions in which shape can evolve (Cheverud, 1982; Goswami et al., 2014; see also Rohner, 2025). So while we may expect some mapping between adaptive peak geometry and phenotypic ellipse geometry, the two are certainly not isomorphic. But even granted this correlation, the SGVH alone predicts only the *volume* of the phenotypic ellipse, not other aspects of its geometry. If, and how, an increase or decrease in phenotypic variance might influence a global integration metric like V_rel_ is not immediately clear (although it can affect the internal covariance structure, Petak et al., 2026). What is clear is that Bookstein is correct in highlighting the importance of the magnitude of phenotypic variance, because there is theoretical and empirical evidence that it varies under more- or less-stringent selective regimes in the way envisioned by Simpson. Bookstein is also correct in his assertion that V_rel_ does not carry information about this quantity, at least not directly, because it is divided out. The magnitude of the phenotypic (or Procrustes) variance is therefore the first parameter considered here (Table 1); it is the trace of **C** (Drake and Klingenberg, 2010), or equivalently the sum of the eigenvalues derived from the reordination of **C**.

**Figure 1.**
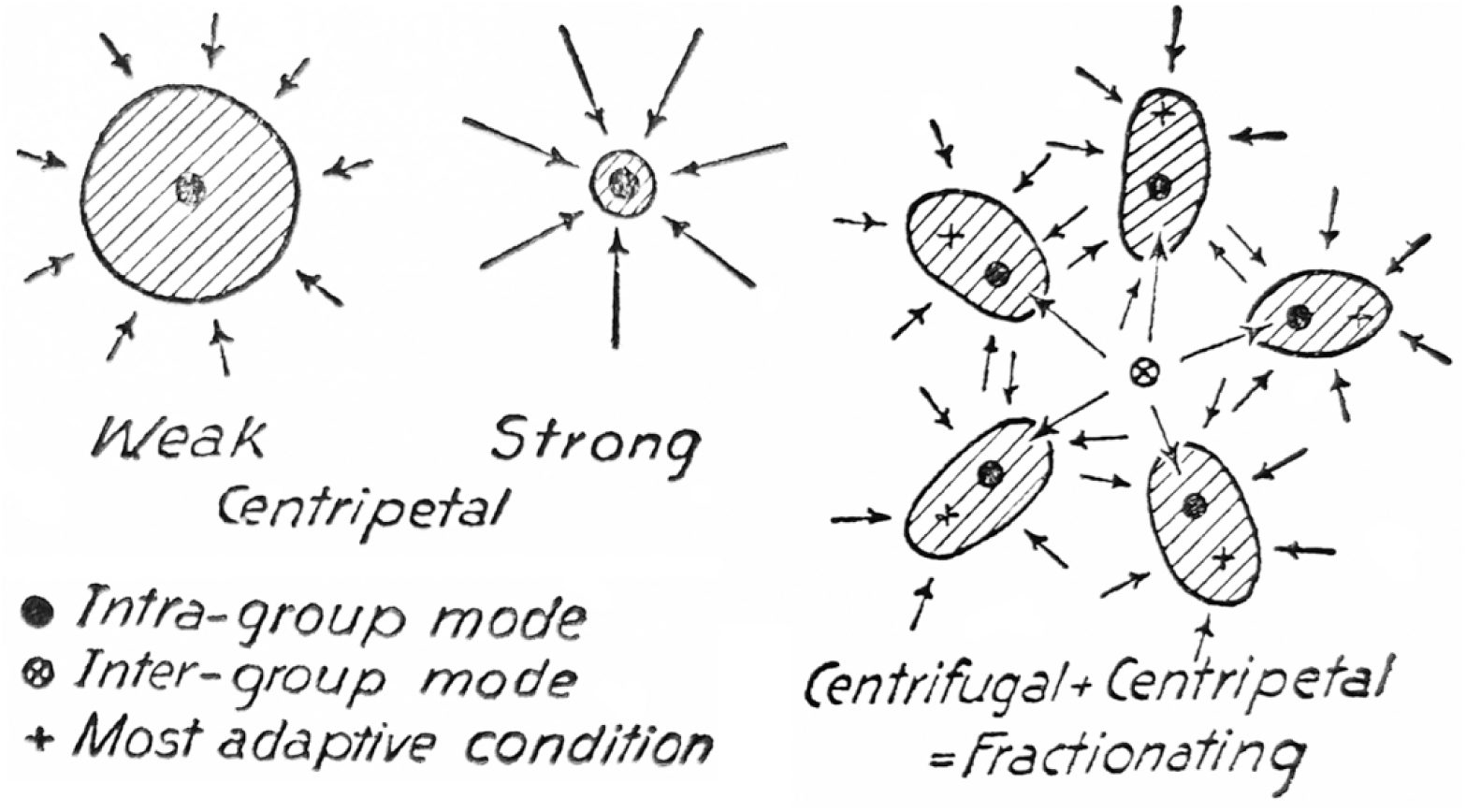
Simpson’s (1953) schematic summary of phenotypic ellipse morphology on a single adaptive peak, with effects predicted for strong and weak selection pressure. Relaxed selection is predicted to result in greater variance. In B, a more rugose landscape is visualized with multiple adaptive peaks separated by local minima. In this case intergroup variance contributes increasingly to the total variance, and the global mean is maladaptive (modified from Simpson, 1953).

## Methods and Material

### Definition of Parameters

The parameters of ellipse volume (sum of variance), dimensional depth (effective rank), and strength of covariance (integration) together allow characterization of the phenotypic ellipse, if relative orientation is disregarded. Strength of covariance is measured here by the rapidity of the eigenvalue decay, because a steeper decay arises from larger early, and smaller late, eigenvalues. Geometric morphometric landmark data generally show a well-behaved, exponential decline, and characterizing this decay directly is desirable (see below). This is done here by first taking the logarithm of the eigenvalues to linearize the decay, and then fitting a simple linear regression to the logged eigenvalues (Results). The slope of this line can then be used to characterize the steepness of the eigenvalue decay. The validity of this exponential decay model for geometric morphometric eigenvalues may be assessed by the R squared of the regression, and these are uniformly high.

Granting for the moment that a log-linear model is appropriate, one can calculate the equivalence of the eigenvalue variance to the slope of the line and the effective rank. Because it is linear, the (ln)eigenvalue distribution is technically a *continuous uniform* distribution. The variance of such a distribution is a simple function of the start and end points for the line, or in this case the first (largest) and last (smallest included) eigenvalues:

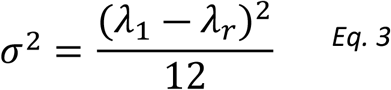

The standard deviation of the eigenvalues is the square root of the above:

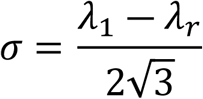

Conveniently, the slope (*S*) of the line modeling the eigenvalue decay can also be expressed in terms of its endpoints, as ‘rise over run’, where the length of the run is the effective rank (*R*), or the number of included eigenvalues:

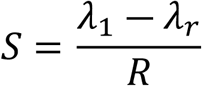

Solving for the eigenvalue difference term and substituting yields:

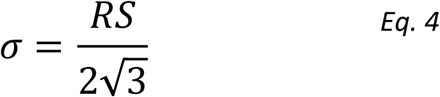

The standard deviation of the eigenvalues is therefore directly proportional to the product of the slope and the rank of the line representing the eigenvalue distribution. The standard deviation is directly proportional to both elements, but in cases where one element increases and the other decreases their product may decrease or increase. The standard deviation is therefore a composite metric of two parameters, and metrics derived from it will be a function of both. In fact V_rel_ is a function of *three* parameters, *R*, *S*, and the phenotypic variance, as can be shown as follows. One may rewrite the equation for V_rel_ given by Watanabe (2022) and shown here in Table 1 as:

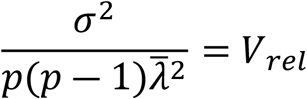

Which in the limit of large *p* reduces to:

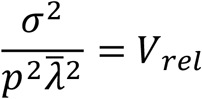

And rewritten as follows:

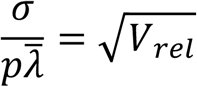

However, the sum of the eigenvalues—the phenotypic variance-- is equivalent to *p* times the mean eigenvalue. Therefore substituting quantities and rearranging yields:

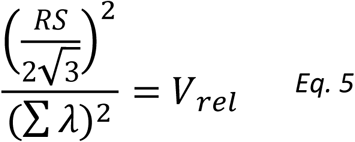

Demonstrating that the relative eigenvalue variance can be rewritten as a function of the three parameters of decay slope, effective rank, and phenotypic variance. The central theme of this paper is that these three parameters need to be considered explicitly, because interpretation of V_rel_ as a summary statistic suffers from their confounding.

Possible impacts of changes in total phenotypic variance on V_rel_ are illustrated in Figure 2. Changes in variance can be accommodated in different ways: if effective rank increases, the slope of the decay decreases and appears more modular (2A); if instead increased variance falls into the first several eigenvectors the decay slope increases and appears more integrated (2B). The slope can also remain stationary, leaving integration unchanged (2C). In the first case V_rel_ is indirectly proportional to the change in variance; in the second it is directly proportional; and in the third it is invariant (given appropriate choice of the eigenvalue cutoff, *X_arb_*). Therefore, in isolation, V_rel_ cannot distinguish between these three cases. An important caveat is shown in Figure 2D; all of these geometries assume that the proportional relationship between variance and matrix information (expressed as effective rank) is constant. If this is true, the decay slope must increase as information decreases, and vice versa, so that the decay slope and information content are dependent. However in real examples this is not the case, as is shown below. Lastly, the three cases in Figure 2 are obviously not mutually exclusive. This paper poses two questions: do closely related species vary when characterized using these three parameters? And if they do, what this mean biologically?

**Figure 2.**
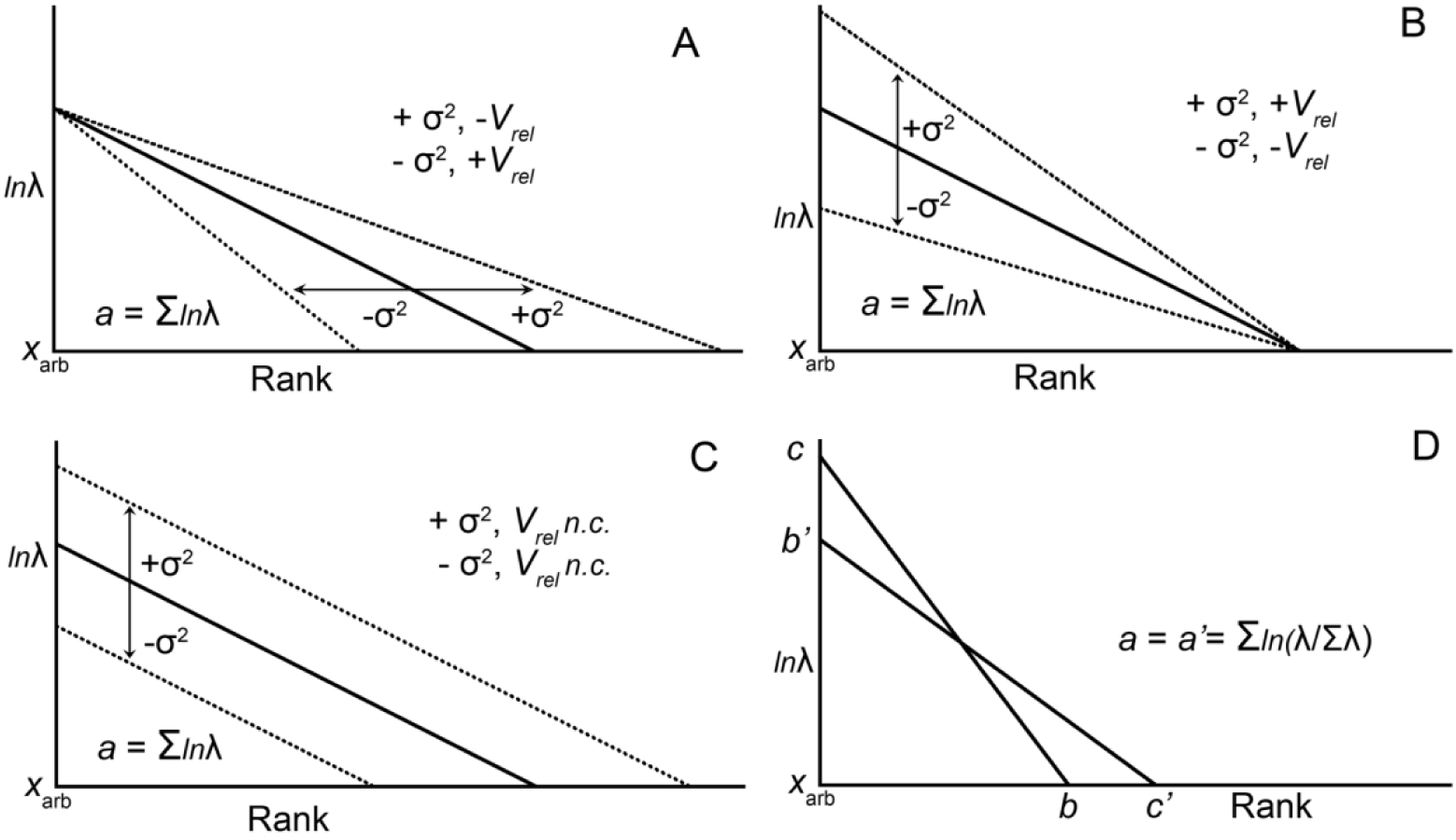
Schematic representations of different cases of how changes in variance can be accommodated in the (ln)scree plots derived from morphometric data. The eigenvalues are modeled as a line, with effective rank (r; the number of included eigenvalues) set at an X value determined by the Shannon entropy. The phenotypic variance (a) will be the sum of the area under the curve. Differences in how increase or decrease in variance is accommodated can have contrasting effects on the variance of the eigenvalue distribution (V_rel_), and the eigenvalue variance is therefore not a unique specifier of any of the cases. For further discussion see text.

### Permutation Test

Another critique made by Bookstein (2022) of single metrics of phenotypic integration concerns the appropriate null model used to scale the metric. In his study, Watanabe (2022) simulates distributions out to the case of the *p*-ball, or complete sphericity. Bookstein argues that this is not realistic, as biological systems are highly correlated. While this is true, it begs the question of what an appropriate null model of covariance might be. O’Keefe et al. (2022) addressed this need by implementing a permutation test on the original data matrix, yielding the quantity of permuted effective rank (πR_e_). This parameter is the average effective rank of a collection of permuted versions of the data, in which each variable vector is randomly resampled without replacement. This will randomize the covariance among variables while maintaining the variance of each variable unchanged. This general approach is similar to that employed by Adams and Collyer (2016), although in that case the variable vectors were drawn from a normal distribution rather than from the original data. A very similar permutation test was employed by Björklund (2019), although in that case the rows were permuted rather than the columns. The permuted effective rank is an important quantity because it gives a measure of the inherent rank deficiency, or amount of information, in the data set without biological covariation (Björklund, 2019; O’Keefe et al., 2022). In **C** derived from a rank deficient, permuted matrix, the off-diagonal covariances will continue to be non-zero due to a fundamental lack of information in the underlying data. It is argued in O’Keefe et al. (2022), and here, that this quantity is an appropriate one with which to compare the sample parameters described above. Again using the *Aenocyon* example from O’Keefe et al. (2022), the permuted effective rank is just over 21 for a real effective rank of 11.32. This is still less than the expected 24, but is greater than the number of landmarks; increasing the sample size should drive the permuted effective rank toward the theoretical maximum of 24 (Conaway and Adams, 2022). A possible methodological issue here is that the permuted matrices no longer reside in the shape space of the original data, and so one may question whether they are appropriate for comparison. They are appropriate because only the rank of each permuted hyperellipse is used for comparison, not the covariance structure within each permuted hyperellipse. The permuted effective rank is a function of the eigenvalues only, not their associated eigenvectors, and therefore the internal covariance structure of the shape space is not relevant.

### Quantities Measured

Once a null expectation for dimensionality has been established using permutation, creating a metric for the amount of dimensionality reduction is straightforward. One may calculate the ratio of the permuted effective rank to the real effective rank:

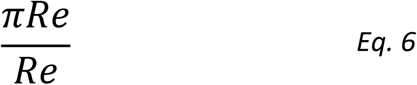

The permuted, ‘uncompressed’ rank is in the numerator, and the compressed rank is the denominator. This is analogous to the general convention for data compression in signal processing contexts, with uncompressed information in the numerator and compressed information in the denominator. Another advantage of this *phenotypic compression ratio* is that it is a dimensionless quantity.

Also, the slope must be compared to a null expectation. The permuted matrix described above can also be utilized to normalize the slope of the eigenvalue decay: slopes from a sample of permuted distributions are first calculated, and the mean of this permuted slope is then used in a ratio with the actual slope:

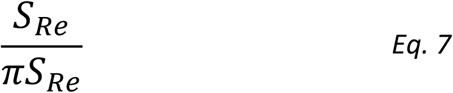

The resulting *phenotypic slope ratio* is normalized to the intrinsic slope of the eigenvalue decay without biological covariance (Table 1). This ratio has a number of desirable properties: it is dimensionless; and it is derived from all relevant eigenvalues via regression.

### Model Systems and Data

The primary data analyzed here are landmark coordinates from the jaws of three closely related sigmodontine rodents (Figure 3). These data originate from Márquez’s (2008) exhaustive study of modularity in a clade of nine rodent species. The taxa used here were chosen to maximize sample size, and to give a range of habitat preference. Of the three species chosen, *Oryzomys palustris* is an ecological generalist with a broad geographic distribution, wide habitat preference, and a varied diet (Rose, 2023). *Nectomys squamipes* is considered a habitat specialist (Ernest, 1986), with webbed feet and a narrow habitat preference, being largely restricted to small stream tributaries (Lima et al., 2016). The third included species, *Microryzomys minutus,* is restricted to higher elevation moist forests, but utilizes a broad range of montaine habitats and is common where it occurs (Carleton, 2015). The phylogenetic relationships of the taxa are known (Figure 3; Márquez, 2008), with *Nectomys* and *Oryzomys* being sister taxa, so that specialist and generalist are each other’s closest relative, with an intermediate taxon as outgroup. The data used here are the landmark data after Procrustes superimposition. The thorough analysis of modularity models by Márquez (2008) also demonstrates that the internal covariance structure of the three taxa is largely conserved.

**Figure 3.**
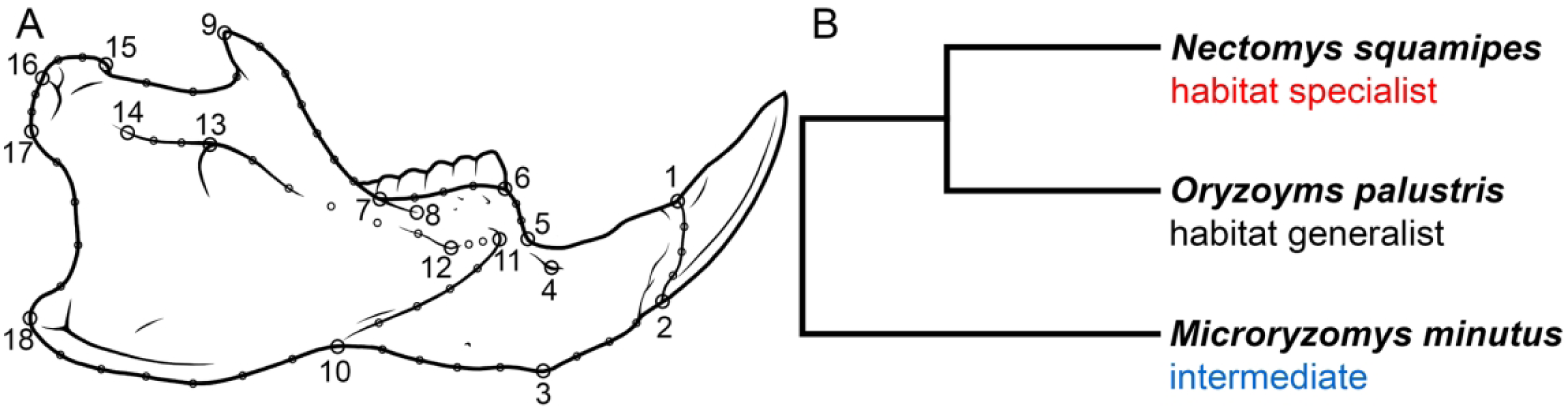
Left lateral view of *Nectomys squamipes*, drawn from Caccavo et al., 2021, with the 18 traditional and 51 semi-landmarks indicated (Márquez, 2008). The phylogeny is a pared version of that in Márquez, 2008, with degree of habitat specialization shown. These assignments are qualitative.

The rat data analyzed here (Supplementary Information 1) are a subset of that analyzed by Márquez, 2008, and was kindly provided by the author. The two dimensional landmarks comprised 18 traditional and 51 semilandmarks, digitized from photographs of right mandibles (Márquez 2008; Figure 3). The provided data were Procrustes-superimposed coordinates and a vector of centroid size, and comprised 546 individuals across nine taxa (six genera; there are three species of *Oryzomys*). From these the three taxa used here were chosen to maximize both sample size and habitat breadth. The sample size for *Microryzomys* was 69, so seven *Nectomys* and one *Oryzomys palustris* were cut for a sample size of 69 in all taxa. The Procrustes variances of the three taxa studied here are calculated as differences from the mean of all nine; this is a benefit because each taxon has a relatively small impact on the mean, so that each taxon is compared to something like a common standard determined by the other eight.

Wolf and dog cranium data were taken from Evin et al. (2025) and comprise 28 three-dimensional landmarks, eight of which are on the midline with ten others mirrored, all digitized from 3D scans of each skull (Supplementary Information 2). Procrustes superimposition of the landmark coordinates for each group was accomplished using the **geomorph** package (*v4.0.10*; Adams et al., 2025) in R 4.1.0 (R Core Team, 2021). The code for the Procrustes superimposition is trivial and is not reported here; complete code for all other analyses performed below may be found in Supplementary Materials 1. The resulting Procrustes coordinates and vectors of centroid size were tabulated and sorted by size. Eighty wolves from the center of the size distribution were taken, while 80 modern domestic dogs were downsampled from the 158 in the data set. The dogs omitted were the seven largest and 71 smallest; this was done to mitigate the much greater size range shown by dogs.

### Analysis

An initial principal components analysis of **C** was performed with the three rat taxa pooled (Figure 4). Inspection of the PCA scores and regression against centroid size shows both high intergroup variance, and a strong association with centroid size (Figure 4A); allometric residuals were therefore calculated by regressing each variable against centroid size (Drake and Klingenberg, 2010; the same procedure was used by Márquez, 2008). All subsequent rat analyses used these allometric residuals (Figures 4, 5). To gain insight into the relative orientation of the point clouds, the method of Common Principal Components was employed (Flury, 1984), implemented in the package **multigroup** in R (Eslami et al., 2020). In this procedure, a single set of common eigenvectors is calculated, and then scores for each group are computed, with only the eigenvalues for each group allowed to vary. The use of intergroup CPCA is discouraged (Houle et al., 2002), so the implementation here is heuristic, but the CPCA vectors are a useful common reference for comparing the relative variance distributions among the three taxa (Figure 4B).

**Figure 4.**
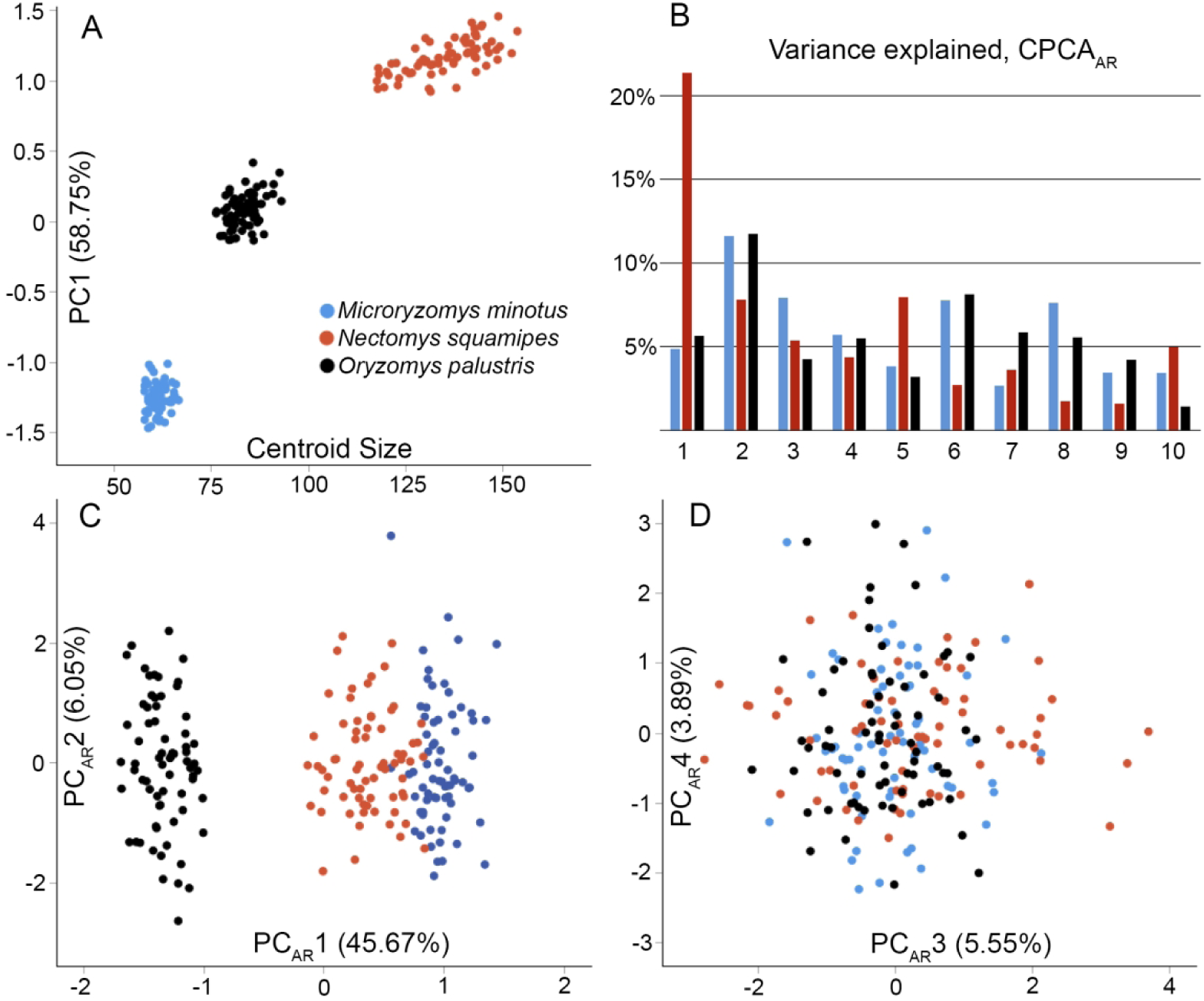
Phenotypic ellipses of sigmodontine rodents. In all plots, *Microryzomy*s is shown in blue, *Nectomys* in red, and *Oryzomy*s in the black. Panel A shows PC 1 from the initial, pooled PCA, plotted against centroid size. PC 1 strongly segregates the taxa, is correlated with centroid size, and contains significant intergroup variance. Subsequent panels use allometric residuals (AR). Taxon-specific loadings for the first ten eigenvectors from the CPCA are plotted in panel B. *Nectomys* loads higher on CPC1 and CPC5, while the other taxa load higher on CPCs 2, 4, 6, and 8; the habitat specialist *Nectomys* concentrates more variance on higher eigenvalues than do the other taxa. The first four PCs (C, D) still recover a PC1 that segregates the taxa, while the others do not. *Oryzomys* differs most in shape relative to the other two. Together these PCs account for 61.16% of the total variance. The range of *Nectomys* on PC 4 is relatively restricted (*p* =0.033, homogeneity of variance test), but is comparable on PC2 and and PC3. Both observations suggest, qualitatively, that *Nectomys* variance is more concentrated on early eigenvalues, and is hence more tightly ‘integrated’.

**Figure 5.**
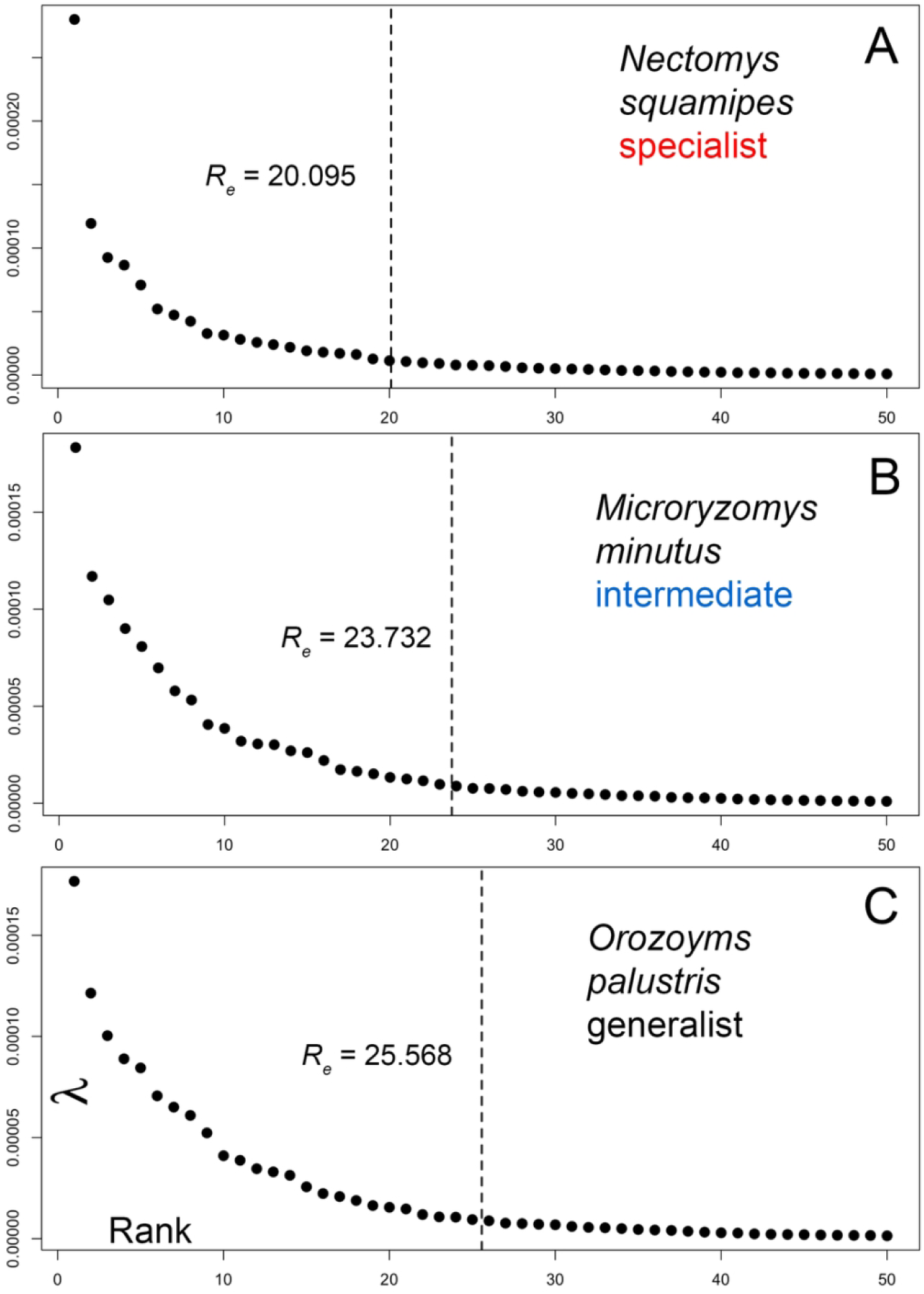
Scree plots showing the eigenvalue distributions of *Nectomys* (A), *Microryzomys* (B), and *Orozoyms* (C), with effective rank (Re) for each shown as a dotted line. The *Orozoyms* matrix contains over five more ranks of information than that of *Nectomys*.

The natural logarithm of the eigenvalue vector was then calculated, and a linear regression fit to the logged eigenvalues. All regressions were performed using the default **lm** tool in R, and the *r*-squared value for each regression is reported within each plot. Only those eigenvalues out to the effective rank of the matrix were used in calculating the regression, with fractional ranks rounded up. Analysis continued by computing the Re and the V_rel_ statistics, using 26 eigenvalues for the rats and 15 eigenvalues for the canids. These cutoffs were chosen as the largest of the effective ranks of the taxa compared. The V_rel_ was also calculated using the appropriate function in **geomorph,** and results were similar to those reported below. In the permutation portion of the code, each column of the original data matrix was permuted, and the effective rank and decay slope of the covariance matrix of the resulting permutation was recorded. This was repeated 5000 times to yield the distributions of the permuted effective rank and permuted decay slopes. Moments of these distributions were then used to calculate the compression ratio and the slope ratio as defined in Table 1, and plotting functions for scree plots and geometry plots are included in the code (Supplementary Information 3). Values for computed parameters may be found in Table 2 (rats) and Table 3 (dogs). Real values are reported in black in Figures 6 and 7, while permuted values are plotted in red; bootstrap estimates and confidence intervals are also in black. Lastly, a version of the code was created that allows for bootstrapping the values in Table 1 (Supplementary Materials 4). This code allows for bootstrapping of confidence intervals (reported below) and subsequent T-tests on parameters of interest, and has the plotting functions removed; these values are also reported in Tables 2 and 3. The tables also report ANOVAs of means and standard deviations, and post-hoc Tukey HSD tests on the means. Because the bootstrap code has a nested permutation loop, bootstrap replicates were held to 1000 bootstrap replicates of 500 permutations each to limit computation time. Importantly, the sample size for the T-tests is the number of eigenvalues included in the eigenvalue distribution, not the number of individuals in the original data matrix; the values used were the bootstrapped effective ranks rounded up (16, 19, and 20 for *Nectomys*, *Microryzomys*, and *Oryzomys*, respectively). For canids, these ranks were 13 for wolves and 9 for dogs.

**Figure 6.**
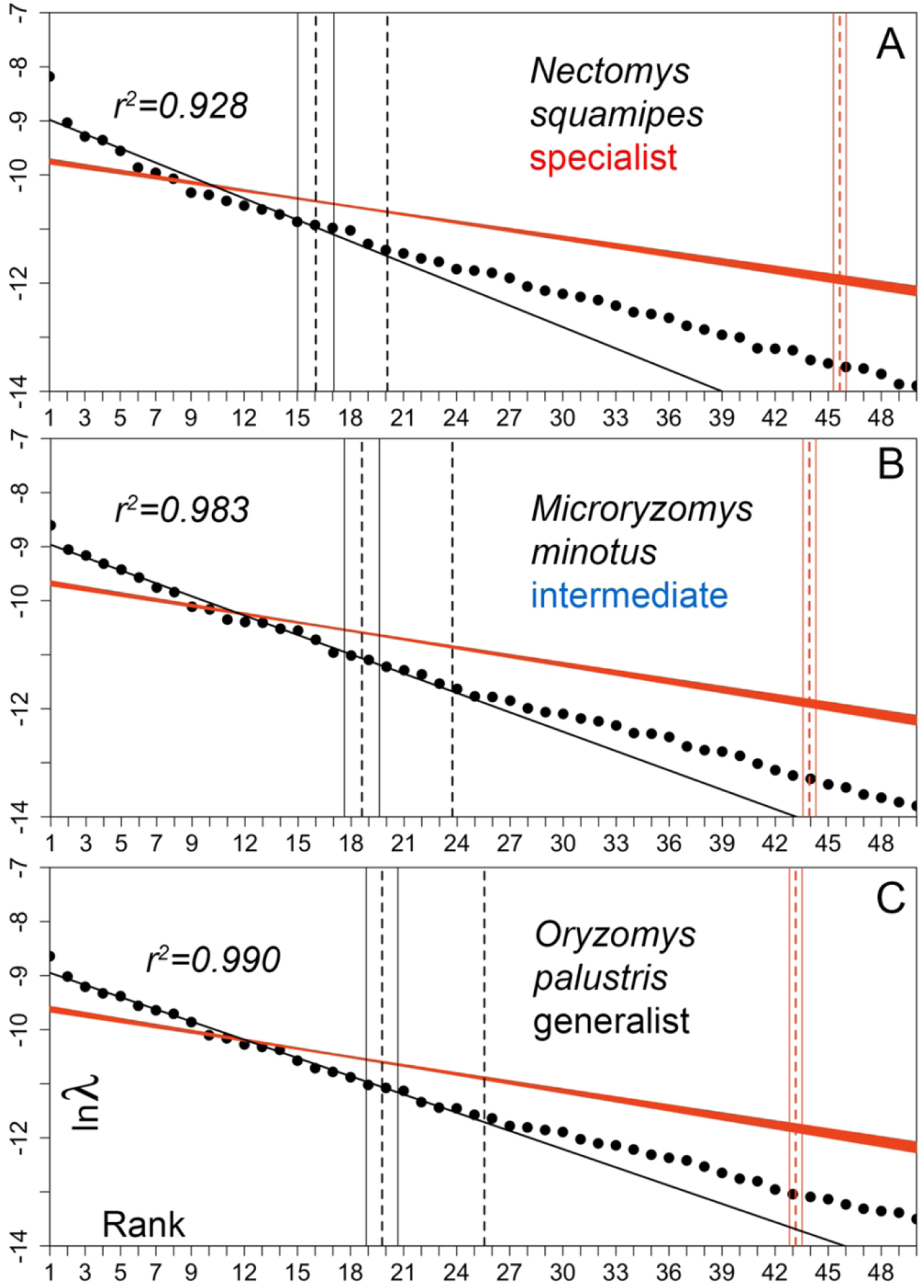
Phenotypic ellipse plots for the three rat taxa analyzed here. Each plot shows rank on the abscissa and the log of the eigenvalue on the ordinate. The solid black line is a regression fitted to the logged eigenvalues, out to the effective rank of the matrix, with fractional ranks rounded up. The red envelope comprises regressions run through the eigenvalues of 5000 column permutations of the original Procrustes coordinate data. The dotted black line depicts the actual effective rank (Re); the dotted red line shows the permuted effective rank (πRe). The bootstrapped effective rank is shown as a dotted line, with solid lines at one standard deviation. Code for calculation and plotting of these plots may be found in Supplementary Materials 1.

**Figure 7.**
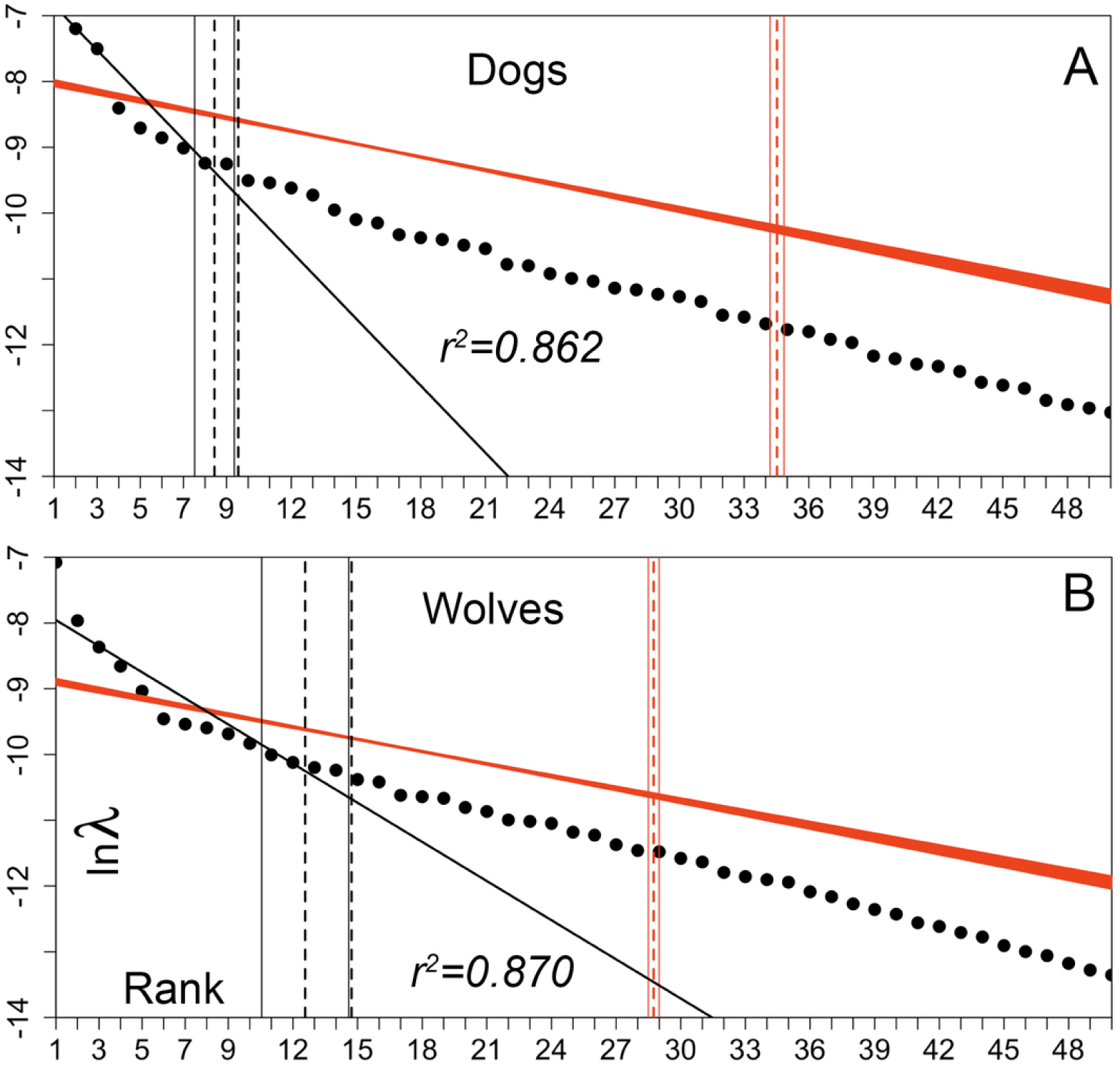
Phenotypic ellipse plots for the two canid analyzed here. Plotting conventions are the same as Figure 6. Dogs are much more tightly integrated than wolves, despite having both much greater overall variance and a higher permuted effective rank. This result replicates that from Drake and Klingenberg, 2010, and is counterintuitive. This pattern may be accounted for by interbreed variance, rather than continuous variance among all dog breeds.

**Table 2.** Calculated phenotypic integration statistics for the three rats in this study. The number of rats in each taxon is 69, with 18 two-dimensional landmarks and 51 semilandmarks (Figure 3). Statistics are: Σλ sum of variance; Re, effective rank; Reπ, permuted effective rank; Cr, compression ratio; Sr, slope ratio; and V_rel26_, V_rel_ calculated for 26 eigenvalues. Sample sizes for T tests and mean comparisons are taken as the bootstrapped effective rank for each taxon, rounded up (16, 19, and 20 for *Nectomys*, *Microryzomys*, and *Oryzomys*, respectively; the permuted effective rank uses the full ranks, or 20, 24, and 26). All statistics are defined in Table 1; code for calculation may be found in Supplementary Materials 1.

| Taxon | *Nectomys* (specialist) |  | *Microryzomys* (inter.) |  | *Oryzomys* (generalist) |  | ANOVA |
| --- | --- | --- | --- | --- | --- | --- | --- |
|  | value | BS est. | Value | BS est. | value | BS est. | P value; T test, means |
| $\Sigma\lambda$ | 0.118 | 0.1163 +/-<br>0.0061 | 0.121 | 0.1193 +/-<br>0.0052 | 0.129 | 0.127 +/-<br>0.0044 | $p < 0.0000$<br>$N \text{ v } M \text{ } p = 0.2163$<br>$N \text{ v } O \text{ } p < 0.0000$<br>$M \text{ v } O \text{ } p = 0.0001$ |
| **Re** | 20.095 | 16.050 +/-<br>1.020 | 23.732 | 18.611 +/-<br>0.986 | 25.568 | 19.783 +/-<br>0.895 | $p < 0.0000$<br>$N \text{ v } M \text{ } p < 0.0000$<br>$N \text{ v } O \text{ } p < 0.0000$<br>$M \text{ v } O \text{ } p = 0.0011$ |
| **Reπ** | 45.653+<br>/- 0.366 | 45.299 +/-<br>0.462 | 43.931+/-<br>0.367 | 43.590 +/-<br>0.547 | 43.151 +/-<br>0.357 | 42.774 +/-<br>0.690 | $p < 0.0000$<br>$N \text{ v } M \text{ } p < 0.0000$<br>$N \text{ v } O \text{ } p = 0.0104$ |
| | | | | | | | $M \vee O \ p < 0.0000$ |
| **Cr** | 2.272 | 2.834 +/-<br>0.178 | 1.851 | 2.349 +/-<br>0.123 | 1.688 | 2.166 +/-<br>0.102 | $p < 0.0000$<br>$N \vee M \ p < 0.0000$<br>$N \vee O \ p < 0.0000$<br>$M \vee O \ p = 0.0003$ |
| **Sr** | 2.705 | 3.370 +/-<br>0.215 | 2.306 | 2.789 +/-<br>0.164 | 2.152 | 2.594 +/-<br>0.126 | $p < 0.0000$<br>$N \vee M \ p < 0.0000$<br>$N \vee O \ p < 0.0000$<br>$M \vee O \ p = 0.0020$ |
| **Vrel<sub>26</sub>** | 0.0688 | 0.0776 +/-<br>0.0123 | 0.0368 | 0.0482 +/-<br>0.0067 | 0.0314 | 0.0422 +/-<br>0.0049 | $p < 0.0000$<br>$N \vee M \ p < 0.0000$<br>$N \vee O \ p < 0.0000$<br>$M \vee O \ p = 0.0361$ |

**Table 3.**
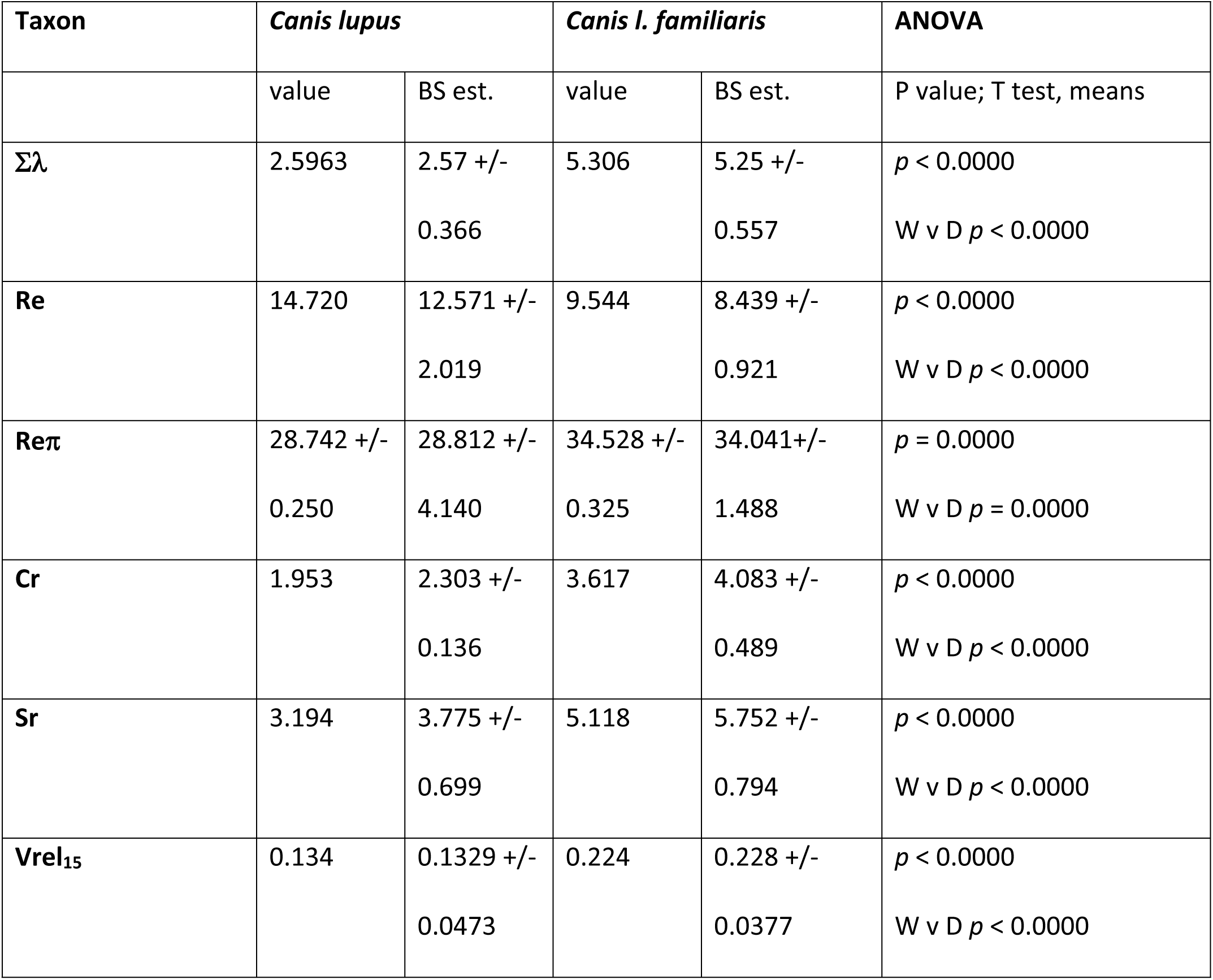
Calculated phenotypic integration statistics for the two *Canis* taxa. Sample size is 80 in both cases, with 8 midline landmarks and 10 mirrored bilateral landmarks. Statistics are: Σλ sum of variance; Re, effective rank; Reπ, permuted effective rank; Cr, compression ratio; Sr, slope ratio; and V_rel15_, V_rel_ calculated for 15 eigenvalues. Sample sizes for T tests and mean comparisons are taken as the bootstrapped effective rank for each taxon, rounded up (13 and 9 for wolves and dogs, respectively; the full ranks were used for the permuted effective rank comparison, these are 15 and 10). All statistics are defined in Table 1; code for calculation may be found in Supplementary Materials 1.

## Results

The first line of Table 2 shows that the phenotypic variances—the volumes of the phenotypic ellipses-- of the three rat taxa differ significantly, with *Nectomys* having the least total variance, and *Oryzomys* the most. Table 2 and Figure 6 demonstrate that the effective rank is also lowest in *Nectomys* (20.095), with *Oryzomys* being over five ranks higher (25.568), while *Microryzoyms* is again intermediate at 23.732. The three parameters are all in agreement, with compression ratio, slope ratio, and V_rel_ all being greatest in *Nectomys* and least in *Oryzomys*. The permuted effective rank shows an interesting pattern; the amount of uncorrelated information is greatest in *Nectomys*, and least in *Oryzomys*. This means that *Nectomys* compresses significantly more information into a smaller phenotypic ellipse, both in volume and dimension, than do the other taxa.

The canid data show that dogs have much greater phenotypic variance than wolves (Table 3), and hence greater ellipse volume, replicating the findings of Drake and Klingenberg (2010). However, the effective rank is greater in wolves despite their diminutive variance, and this difference is large, at over five ranks (Figure 7). The three integration statistics all demonstrate that dogs are much more tightly integrated than wolves, again replicating the findings of Drake and Klingenberg. The permuted effective rank is larger in dogs, meaning there is more total information in the dog data relative to the wolf data. The way this information is compressed is discussed below.

## Discussion

The three rat jaw matrices analyzed here are homogenous by design, having the same number of individuals, the same landmark scheme, and gross differences in body size factored out via the use of allometric residuals. Given this homogeneity, the differences in phenotypic integration parameters displayed by the three taxa are striking. The simplest conclusion one can draw is that *Nectomys* has the least phenotypic variance, and hence the smallest ellipse volume, followed by *Microryzomys*, and then *Oryzomys*. This matches Simpson’s expectations for what we should observe in specialist, intermediate, and generalist taxa, respectively (1953; Figure 1A), and the phylogenetic relations of the taxa prevent these differences from being attributable to phylogeny alone. Obviously the environmental assignments used here are qualitative, and the pattern would be more compelling if niche width were quantified more rigorously (Roughgarden; 1972; Ackerman and Doebeli, 2004; Bearhop et al., 2004), yet the magnitudes of the phenotypic variances do match expectations from the habitat categories. The V_rel_ statistic captures this difference readily, with *Nectomys* having a much higher V_rel_ than the other two taxa, indicating the *Nectomys* is more tightly ‘integrated’ than the others. However, it is not clear from V_rel_ alone where this tightness comes from: whether it is a function of tighter covariance on the significant axes, or a function of the lower dimensionality of the *Nectomys* ellipse, or both. As is clear from Figure 6, *Nectomys* is in fact more tightly integrated in both covariation (the eigenvalue decay is steep) and in dimension (the effective rank is low). Therefore we can characterize the *Nectomys* phenotypic ellipse as being both narrower and shallower than those of the more generalized taxa. This follows from the observation that *Nectomys* has the least total variance, because if there is less variance volume to distribute the ellipse must be smaller. One might argue that the increased V_rel_, along with the smaller phenotypic variance, adequately capture the global increase in ‘integration’ seen in *Nectomys*, because the slope ratio and compresson ratio may simply reflect the paucity of variance in that taxon, and hence are superfluous. However V_rel_ and phenotypic variance in isolation cannot distinguish between the two cases in Figures 2A and 2B. This differentiation requires knowledge of the effective rank.

Effective rank would not be an important parameter if it did not vary so much, being over 5 ranks, or more than 20%, greater in *Oryzoyms* than in *Nectomys*. This can have have a large, and undesirable, effect on the caculation of V_rel_. The V_rel_ statistic was calculated here (for rats) at a constant dimension of 26 eigenvalues (the largest of the three effective ranks, Table 2). However this value arbitrary for *Microryzoyms* and *Nectomys*, because their effective ranks are less than 26, and in isolation we would calculate V_rel_ using fewer axes. Subsidiary analysis shows that when the dimension of each ellipse is allowed to vary with its effective rank, the V_rel_ statistic derived from each no longer captures the differences in tightness shown in Figure 6, indicating that difference in dimensionality is contributing heavily to V_rel_. Therefore calculation of V_rel_ is sensitive to the number of eigenvalues included in its calculation, and comparisons using V_rel_ should include the same number of eigenvalues for each group compared. In fact the rat ellipses analyzed here seem to vary more in dimension than they do in strength of covariance, and it is this dimensional difference that V_rel_ is largely reflecting. In more general terms, V_rel_ confounds the tightness of covariance with the tightness of dimensionality, which may account for the difficulty in pinning down what ‘integration’, as measured by V_rel_, actually means biologically. This confounding is also generally true for all single-value integration statistics (i.e. O’Keefe et al., 2022), which might explain why they have been found not to work (Conaway and Adams, 2022). The advantage of the plots and associated analyses in Figure 6 is that they allow visualization of covariance tightness (the steepness of the eigenvalue decay), dimensional tightness (the effective rank), and phenotypic covariance (the area under the curve). As shown in Equation 5, the V_rel_ statistic is a function of all three of these parameters. Thus far in this paper, the two aspects of VanValen’s tightness—eigenvalue decay and dimensional compression—have been treated as separate entities. Analyzing the two quantities separately is important because they can confound each other (Equation 4). However, it is clear from first principles that the two phenomena must be correlated (Figure 2D). If one holds the amount of variance equal, the eigenvalues must decay more steeply in a matrix with lower rank, and less steeply in a matrix with greater rank. The decay slope is therefore dependent on the effective rank, to an unknown degree. Quantifying the degree of this dependence, and possibly accounting for it, are topics that are beyond the scope of this paper. A simulation study that tracks how the decay slope and effective rank change in concert given a known interior covariance structure is an obvious first step. Further, examination of the dog data presented here demonstrate that tightness in covariance and tightness in dimensionality are not always correlated with the magnitude of the phenotypic variance, meaning the slope decay and the effective rank are at least partially free parameters.

The canid analyses (Table 3; Figure 7) presented here compare 80 wolves and 80 modern dogs with data taken from Evin et al. (2025). These are different crania, with a different landmark scheme, than those in Drake and Klingenberg (2010), but the findings replicate closely the findings of the earlier study. Domestic dogs have more than twice the phenotypic variance as wolves, so their phenotypic ellipse has much greater volume (Table 3). We might therefore expect, given the arguments in Figure 2C, that dogs would exhibit a greater effective rank; in this case the slope decay would be constant and additional variance accommodated by moving the curve up, increasing the rank. However this is not the case; dogs are in fact much more tightly integrated than wolves in dimension, and also in slope ratio. The dog phenotypic ellipse is much shallower than that of wolves, but is also very much wider, with the greater dog variance accommodated on the first few eigenvalues. This is observable grossly using the classic rule of thumb for eigenvalue significance that places the cutoff at kinks in the scree plot. Using this convention, dogs have two significant eigenvalues while wolves have five (Figure 7). Therefore the wolf phenotypic ellipse is both much narrower and much deeper than that of dogs, with much smaller volume, all facts that are readily apparent from Figure 7. The V_rel_ statistic does capture the increased tightness in dog ‘integration’ (Table 3), but the full picture that the dog ellipse is shallower in dimension but much wider on those shallow axes is lost.

The observed pattern in dogs—an increase in dimensional and covariance tightness despite an increase in overall variance— is puzzling on the surface. None of the cases shown in Figure 2 embody the pattern, although there are similarities with case 2A, where additional variance is accommodated by increase in slope. Yet the effective rank decreases markedly, from 15 in wolves to 10 in dogs. This is difficult to achieve given the much higher variance in dogs, at least given a simple fitness surface with a single peak, and so begs a biological explanation. One possibility is that the dog fitness surface, and hence the dog sample, is not homogenous, but instead looks more like Simpson’s more complex adaptive landscape in Figure 1B. A shallow, but rugose fitness surface with multiple steep peaks centered on different dog breeds would yield a range of tightly integrated phenotypes scattered across a broader landscape. This more complex model could yield tightly integrated dogs but with greater overall variance, because intergroup variance will become important. This model is testable using cluster analysis of dog breeds (Sokal and Sneath, 1963), or by simply comparing the intrabreed to interbreed variances within a large sample of dog breeds; these inquires are beyond the scope of this paper. The dog example does demonstrate that V_rel_, in isolation, does not contain enough information to fully characterize phenotypic ellipse geometry. Knowledge of the phenotypic variance and effective rank is also necessary, while separating a single integration statistic (like V_rel_) into dimensional (compression ratio) and covariance (slope ratio) components is also advisable given that single value metrics confound these phenomena.

An unexpected result presented here is that, in both rat and dog species, the permuted effective ranks differs significantly. The permuted effective rank is always much greater than the real rank, demonstrating (once again) that biological shapes are tightly correlated; however the fact that the intrinsic information content should vary between closely related taxa with equivalent landmark schemes and sample sizes is surprising. Of the rat taxa, the specialist *Nectomys* has the greatest permuted rank, and hence the greatest magnitude of information, despite the facts that it has the least phenotypic variance and is the mostly tightly integrated in both slope and dimension. *Nectomys* therefore encodes more information into a smaller phenotypic ellipse than the other taxa, and is hence more efficient in information theoretic terms. The generalist *Oryzoyms* has the least intrinsic information but the largest and most globose phenotypic ellipse, and therefore is less efficient at compressing its information. Caution is advisable here because the ranks involved are high, on the order of 45, and a sample size of 69 is not adequate to make inferences about this many parameters (for discussion see Pavlicev et al., 2009b). However, it is intriguing to note that the specialist taxon appears to encode more information, and does so more efficiently, than does the generalist taxon. Does this reflect more stringent selection on a narrow, steeper adaptive peak? If so, does this impose more stringent constraints on the amount of information a species must carry, and how efficiently it is encoded? These questions are left for further study. The canid example is also illustrative; dogs have much greater permuted information than do wolves. This pattern is similar to that seen in rats, with the more tightly integrated taxon having greater information, and lends credence to the idea that dogs are in fact tightly adapted to a rugose surface of local peaks centered on different breeds. However it this is true, both the phenotypic variance and the permuted effective rank will have unknown, and possibly large, contributions from interbreed variance. Simpson’s simple model of a single fitness peak is no longer appropriate, and characterization of a single dog phenotypic ellipse is probably not tenable.

## Conclusion

This paper explores analysis techniques designed to characterize the geometry of the phenotypic hyperellipse. Minimally, full characterization requires the phenotypic variance, the effective rank, and a measure of covariance strength (like V_rel_ or the decay slope). A single-valued integration metric like V_rel_ is shown to be unable to discriminate between different ellipse shapes, validating Bookstein’s indictment of its use, at least in isolation. These metrics are also vulnerable to differences in the number of eigenvalues used to calculate them, and important issue given the large ranges in dimensionality shown by the rat and dog taxa. Simpson’s (1953) qualitative model of a phenotypic ellipse on a fitness surface predicts that more specialized taxa should be less variable than generalist taxa. This is observed, with the habitat specialist *Nectomys* having less phenotypic variance than *Oryzoyms*. *Nectomys* is also more tightly integrated in both covariance and dimension, having a steeper decay slope and lower effective rank than *Oryzoyms*. These findings lend credence to the interpretation that the *Nectomys* adaptive peak is steeper, and that the geometry of its phenotypic ellipse has evolved to reflect this. The canid example is more nuanced; dogs have much more variance than wolves, and yet are more tightly integrated in both decay slope and dimension. This may be attributable to the dog adaptive surface being a rugose landscape of steep adaptive peaks centered on different breeds: a testable hypothesis. Lastly, the rat model is used to examine the compression of information into the phenotypic ellipse. The specialist *Nectomys* is found to encode information most efficiently (most compression), while *Oryzomys* is least efficient (less compression).

## Supporting information

Data Set 1

Data Set 2

## Acknowledgments.

This paper has benefitted greatly by discussion with M. Zelditch, P. D. Polly, M. Conaway, and J. Spear, as well as extensive feedback from three anonymous reviewers. Special thanks to J. J. Sepkoski for inculcating a healthy distrust of ratios.

## Data and materials availability

All of the data employed in this paper are available in Supplementary Data Table 1 (Rats) and Table 2 (Dogs). The rat data comprise landmark coordinates after Procrustes superimposition, along with a vector of centroid size, and are a subset of those presented in Márquez, 2008. The dog data are raw landmark coordinates, and are a subset of those presented in Evin et al., 2025; for further details see the source publications. All R code used in this paper is available in the Supplementary Information.

## PermTestPlot.R

okeefef

2026-02-02

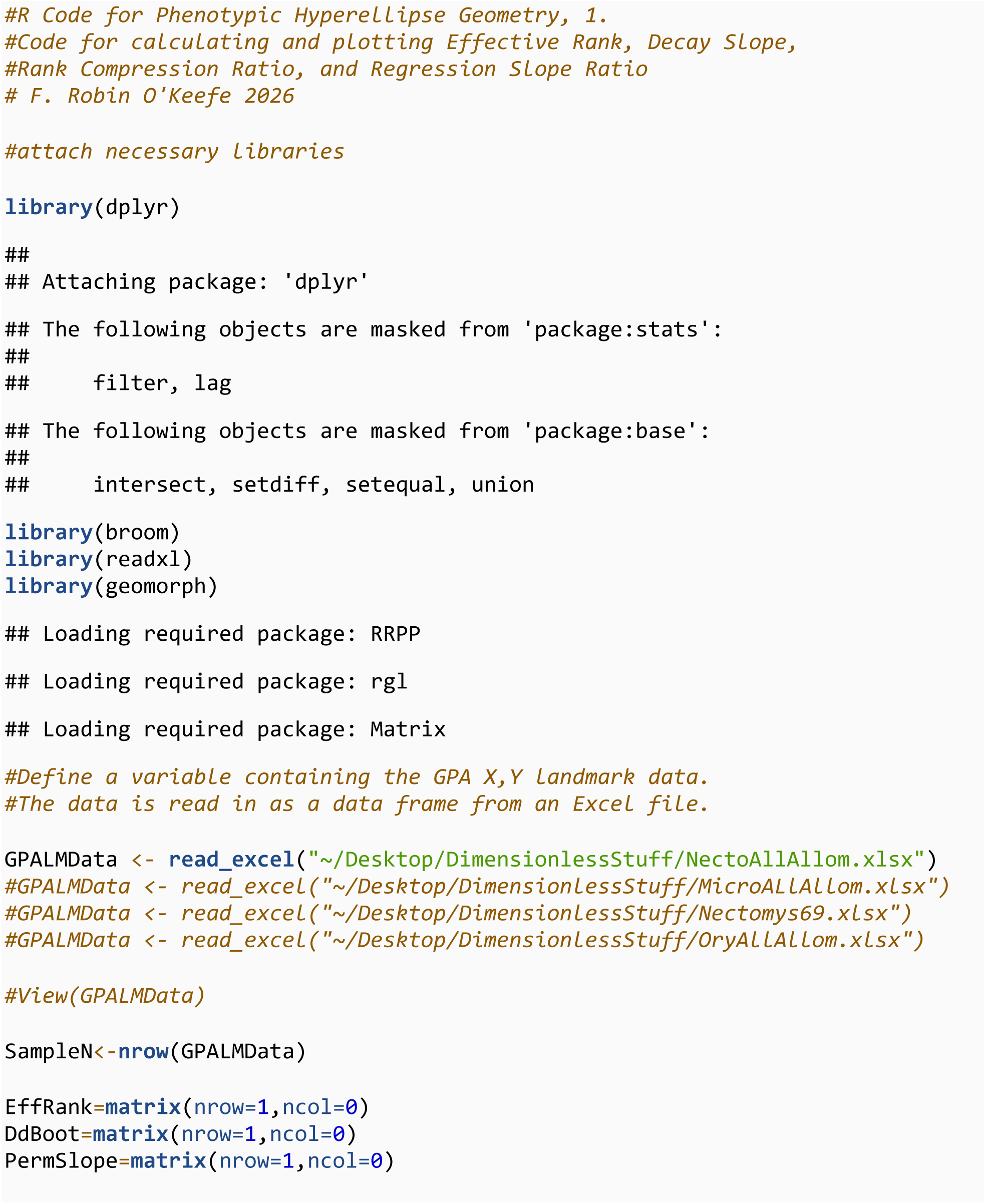
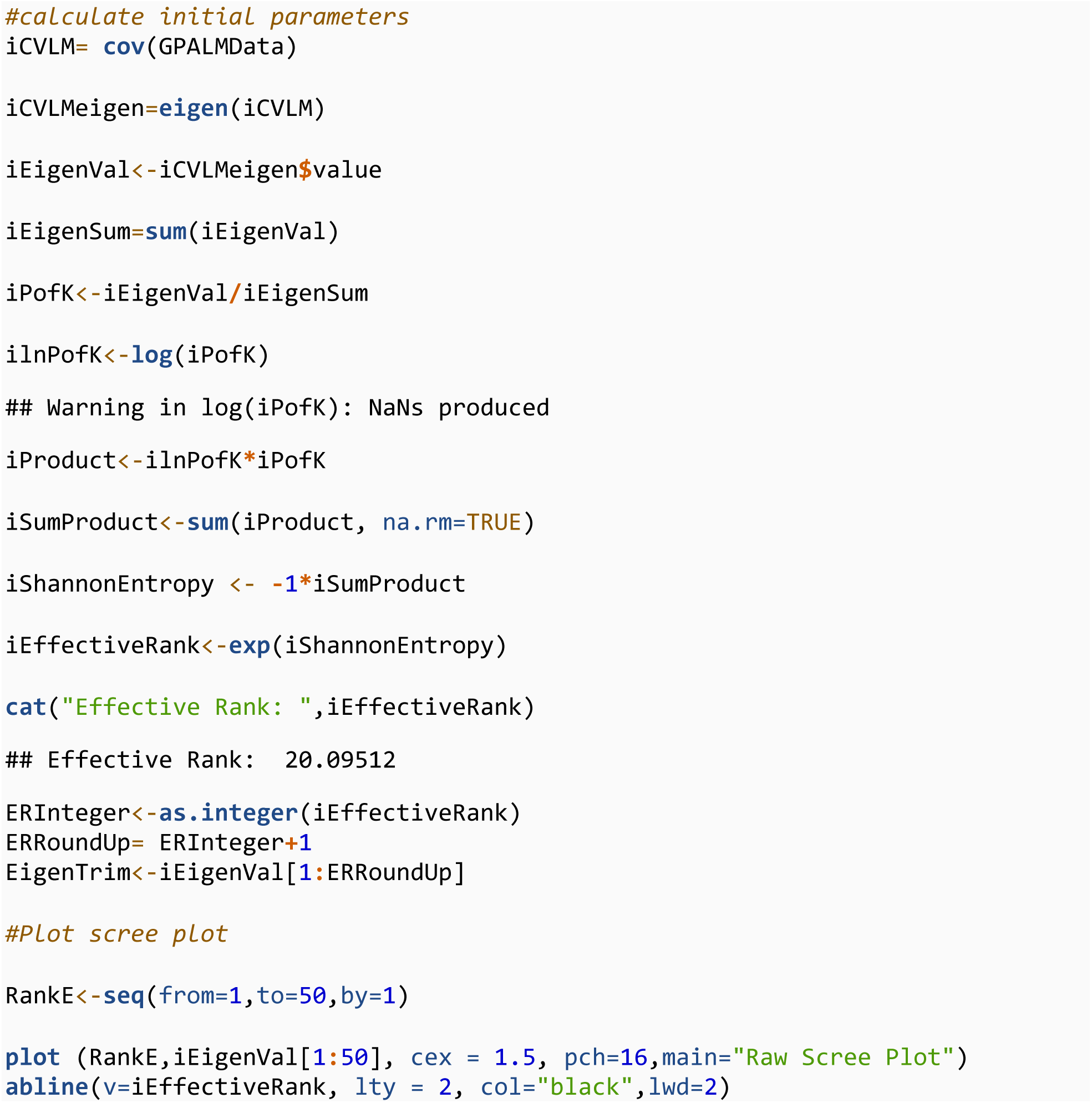
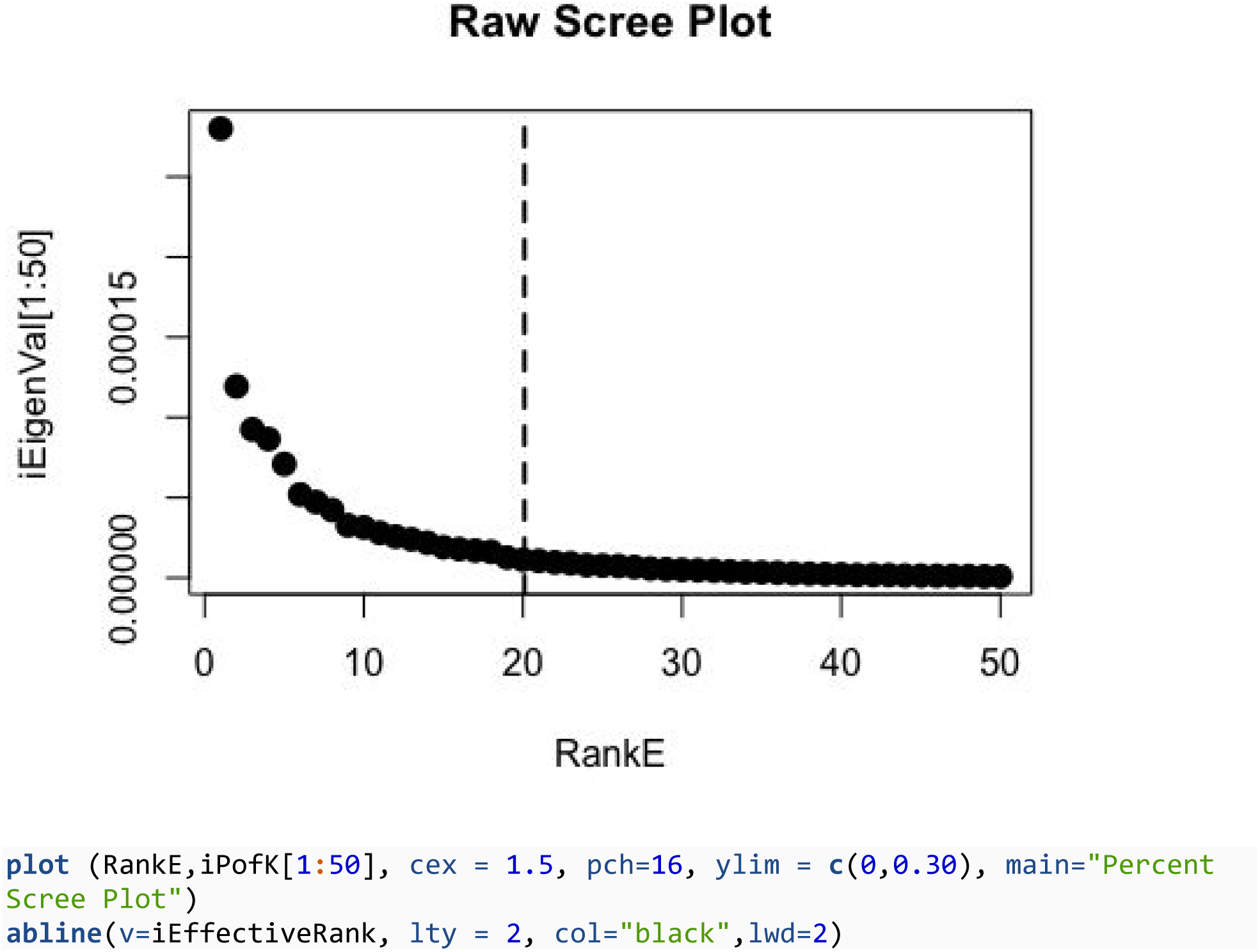
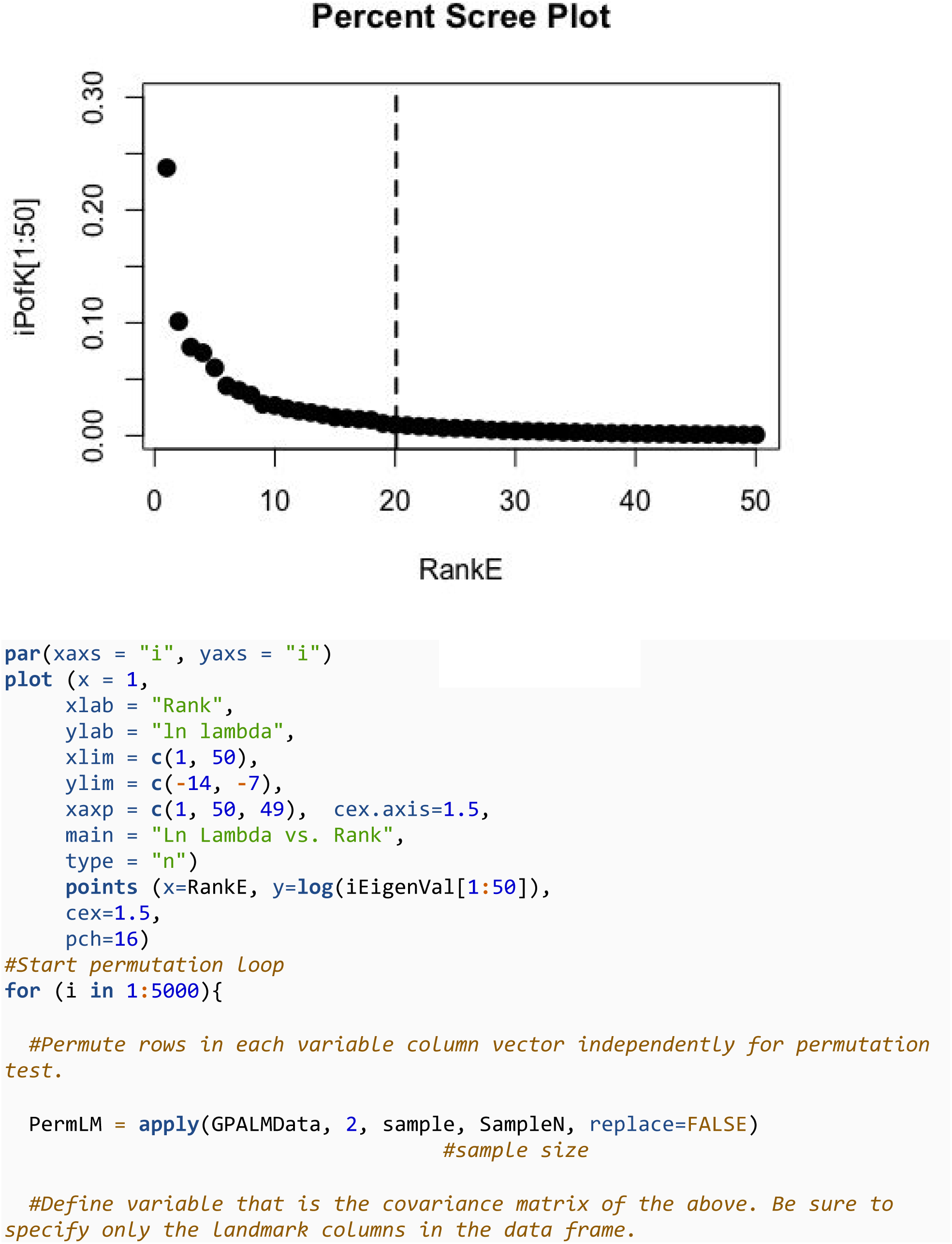
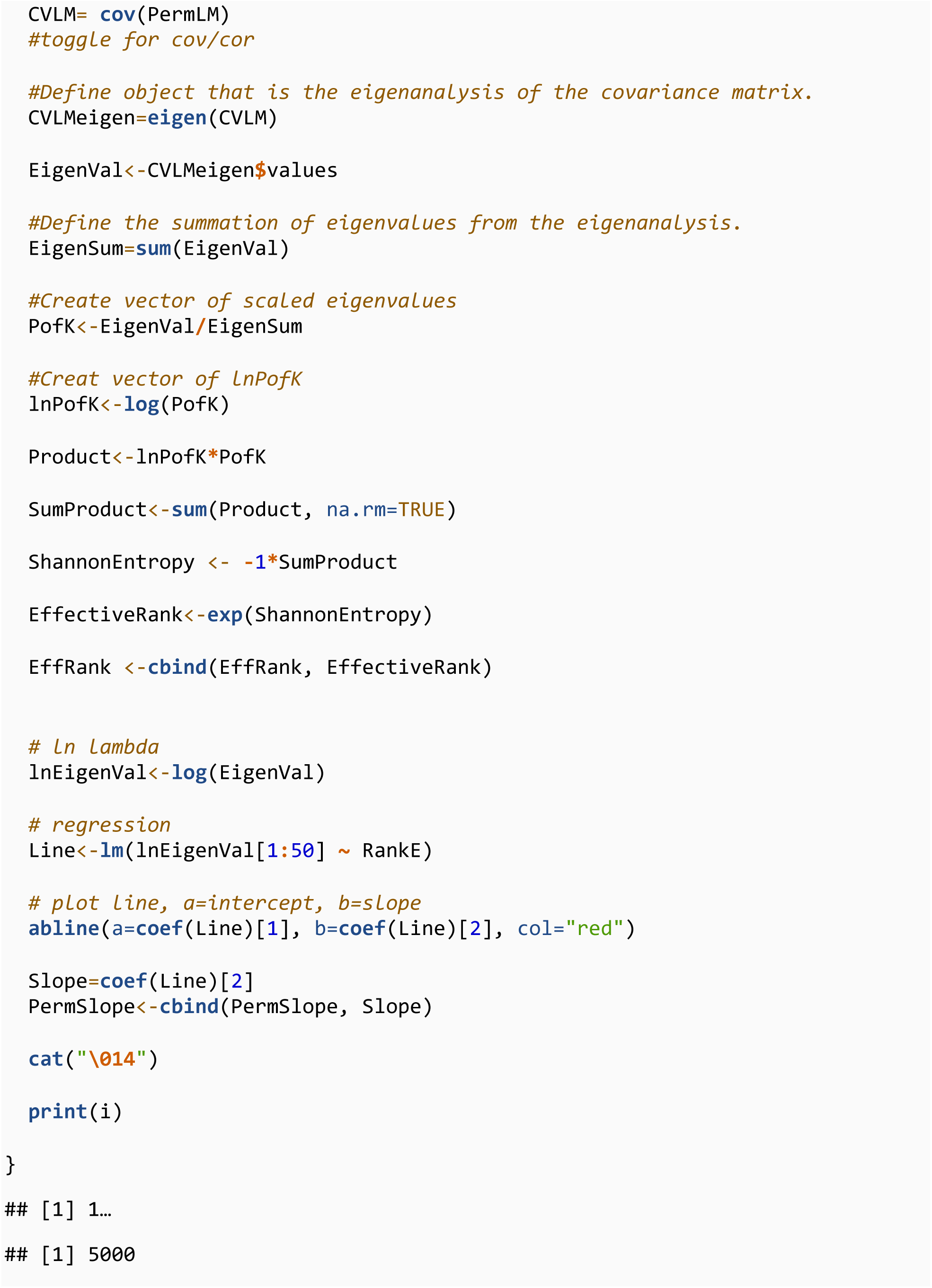
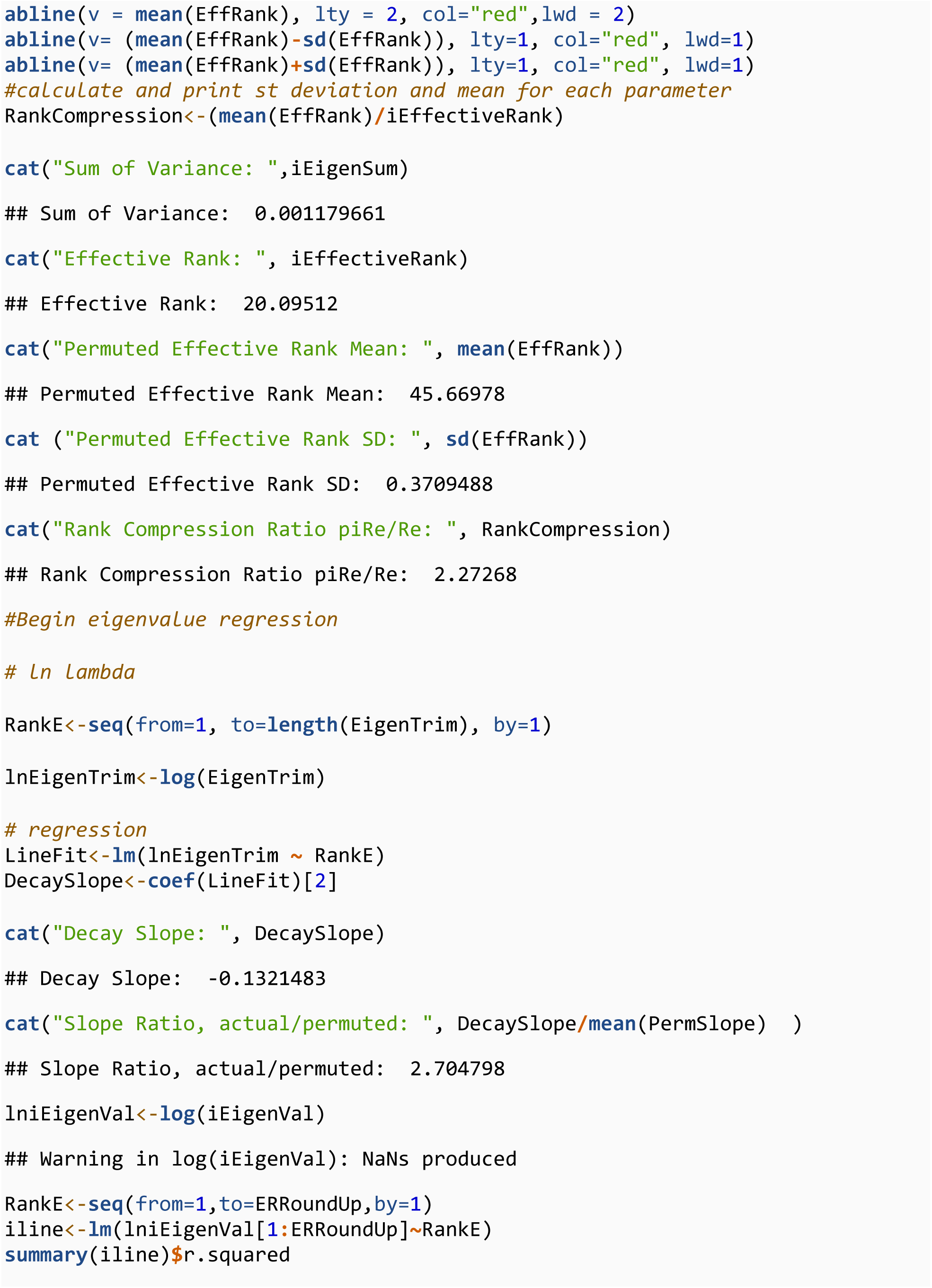
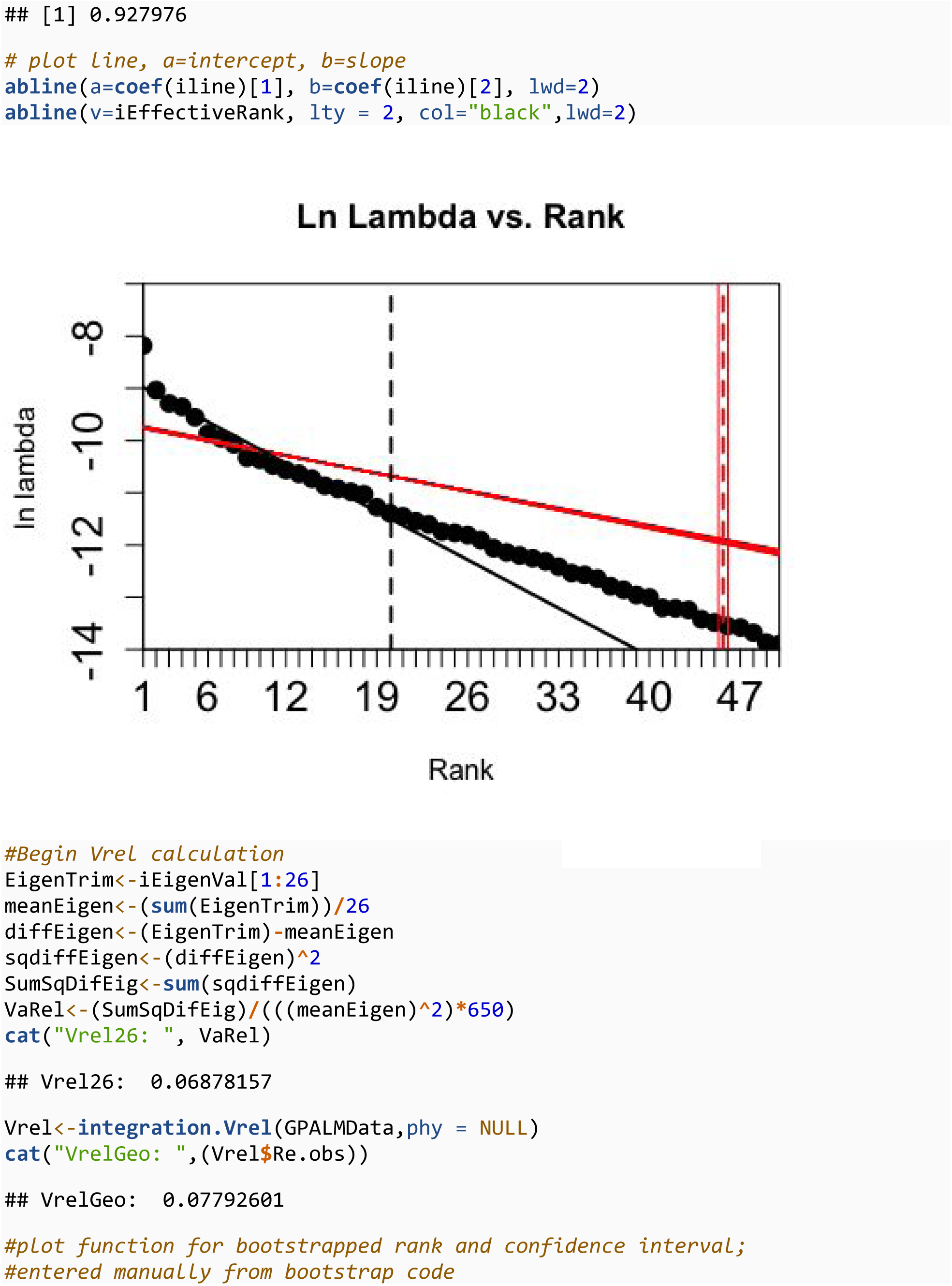
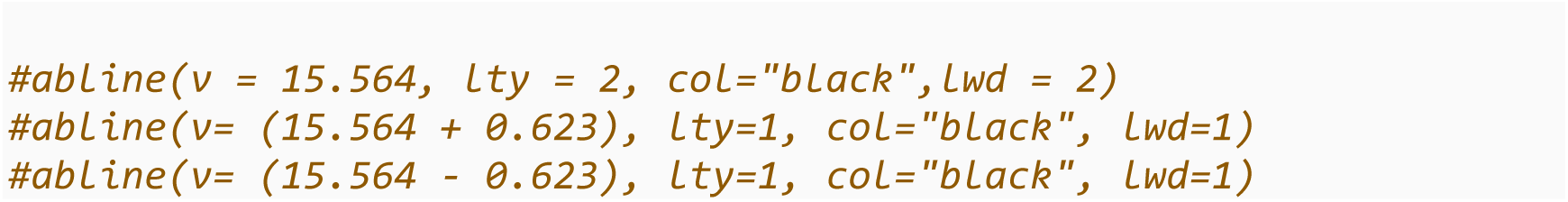

## PermTestBoot.R

okeefef

2026-02-02

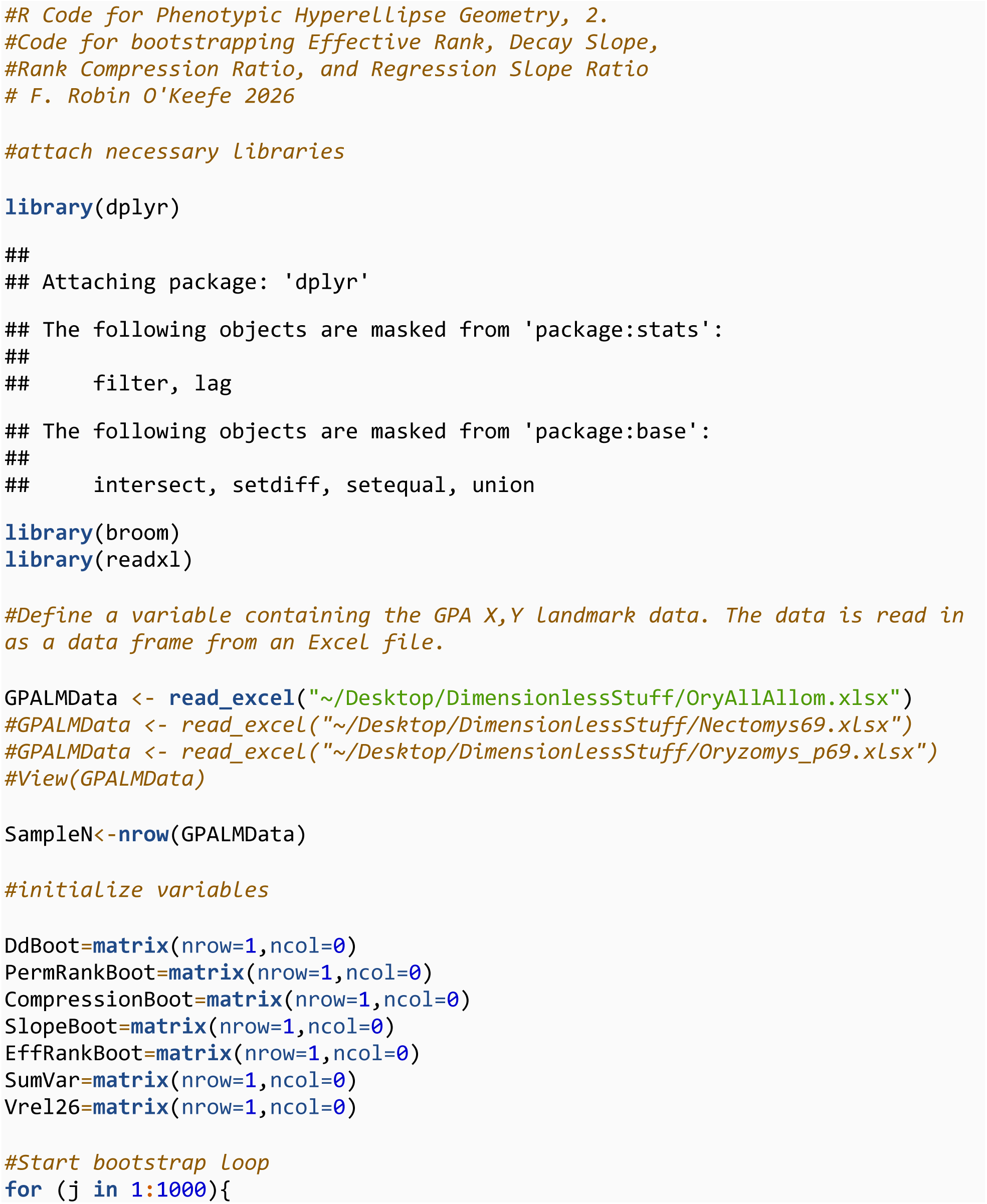
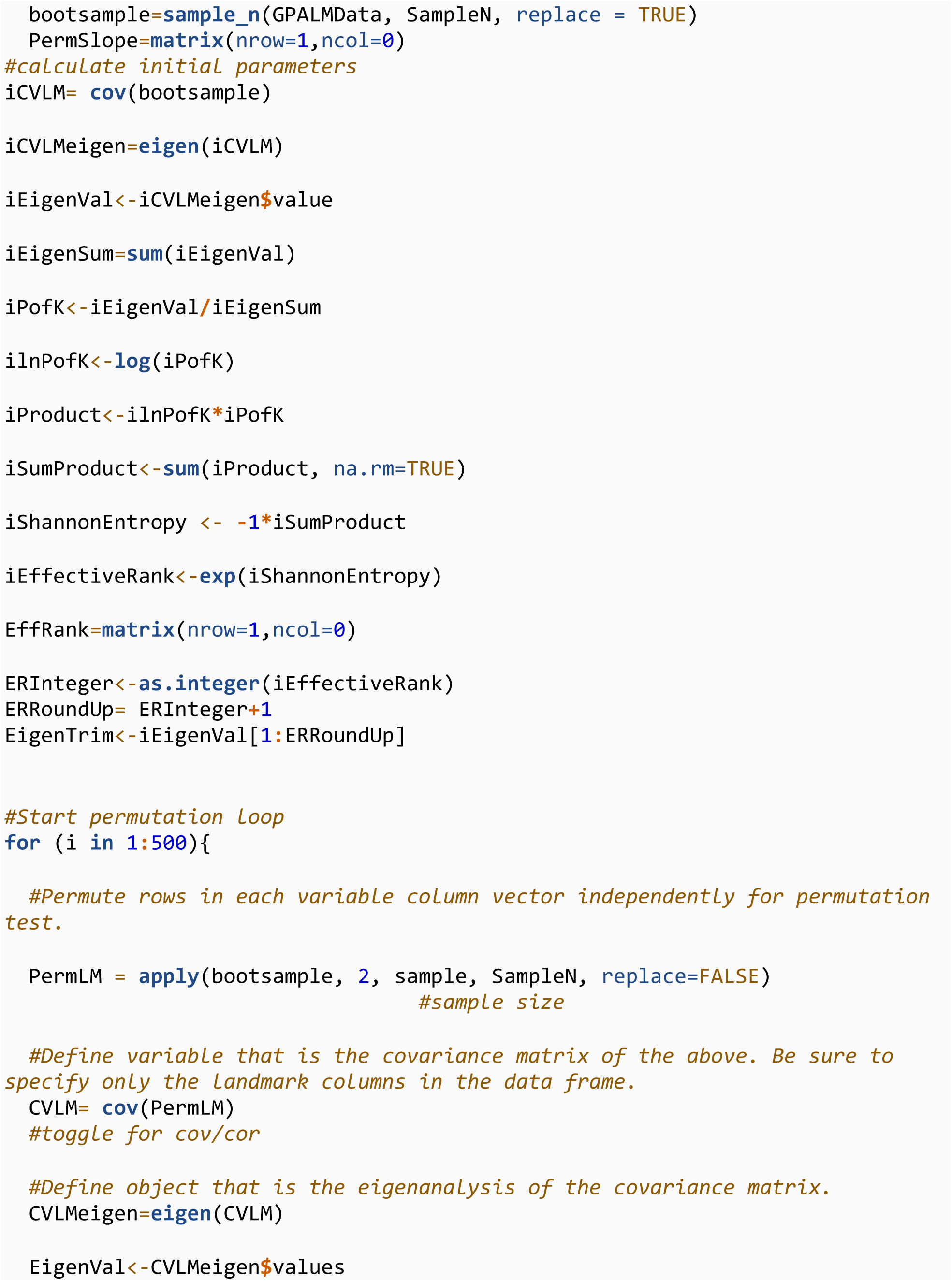
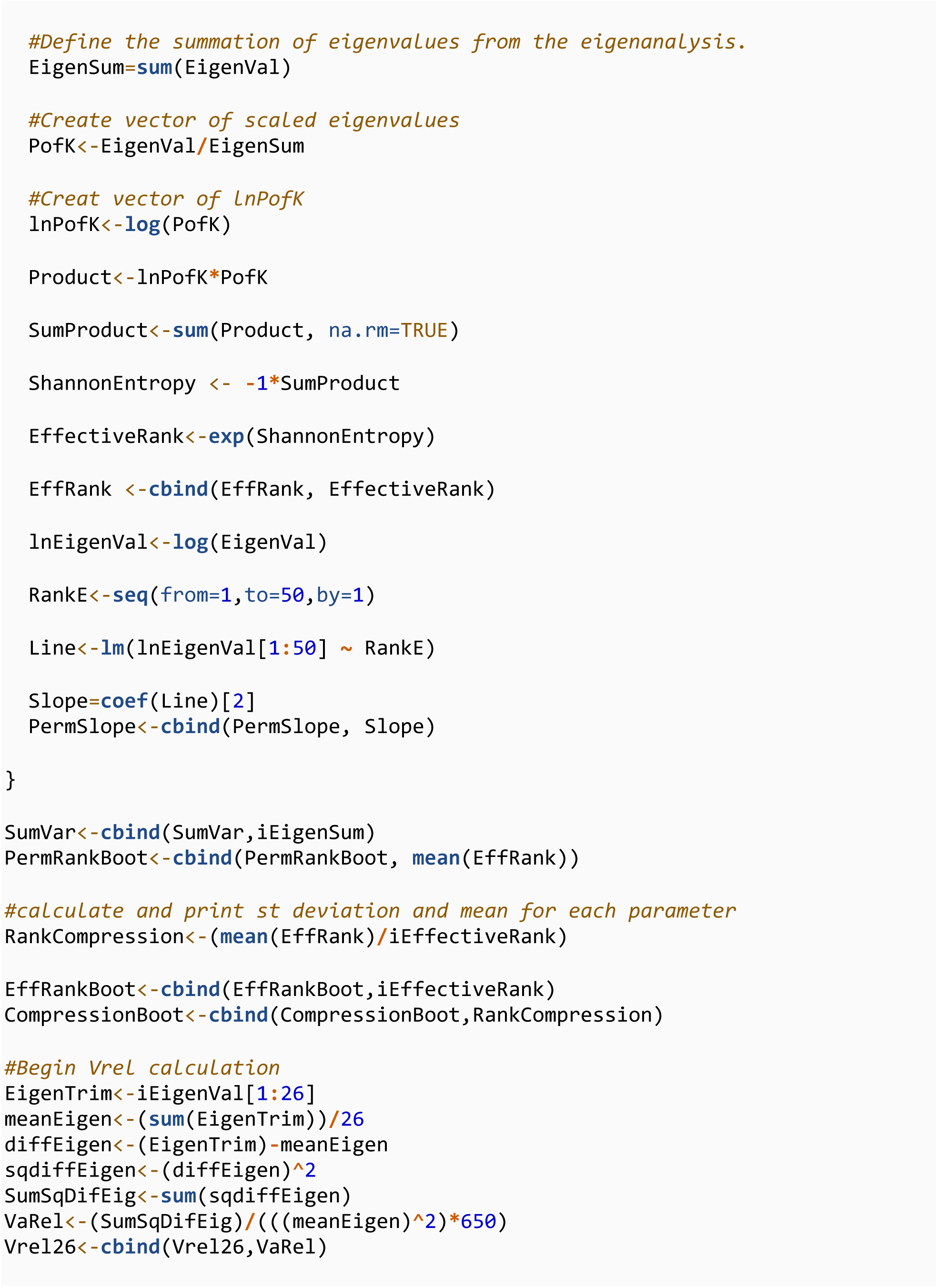
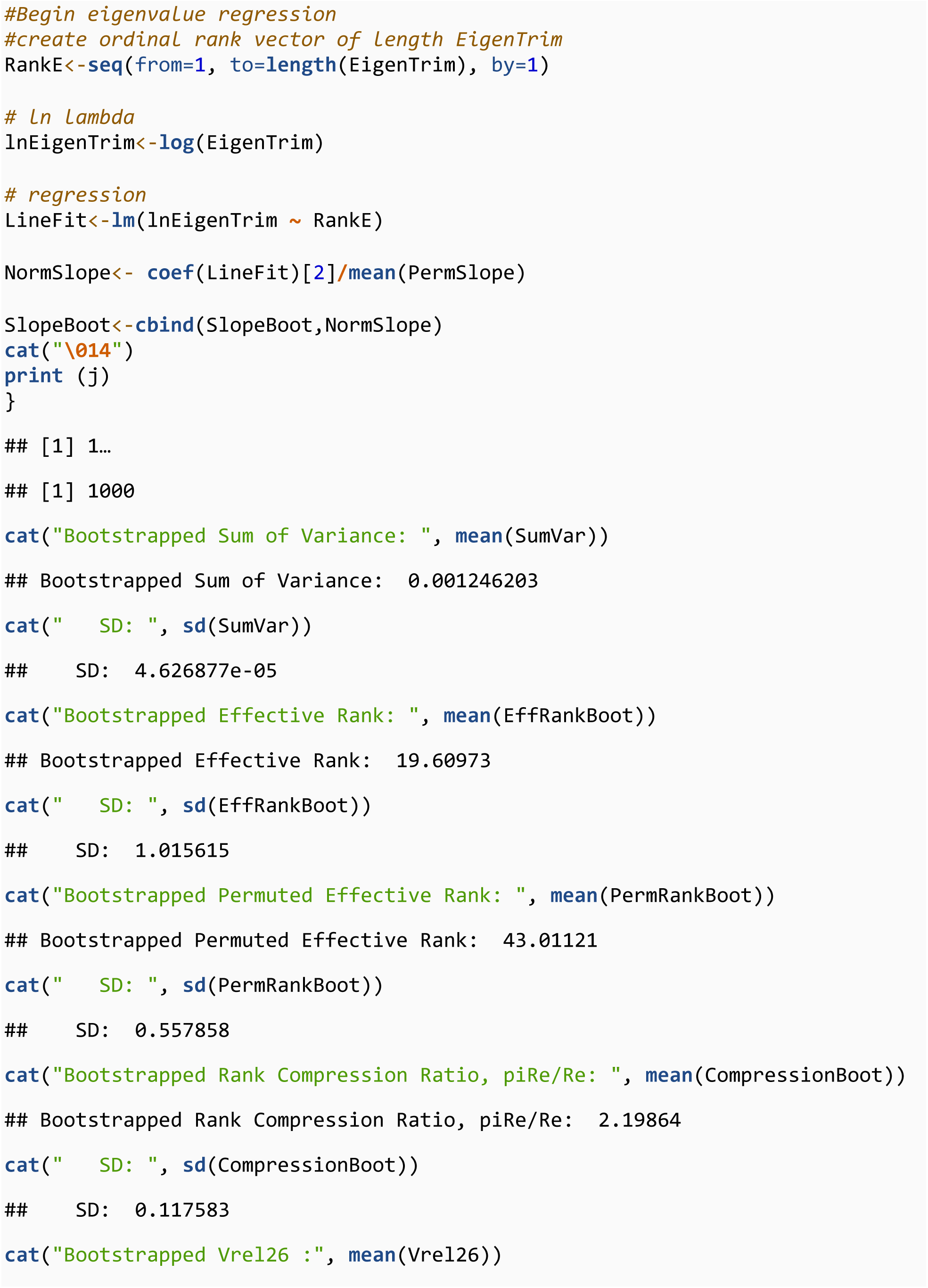
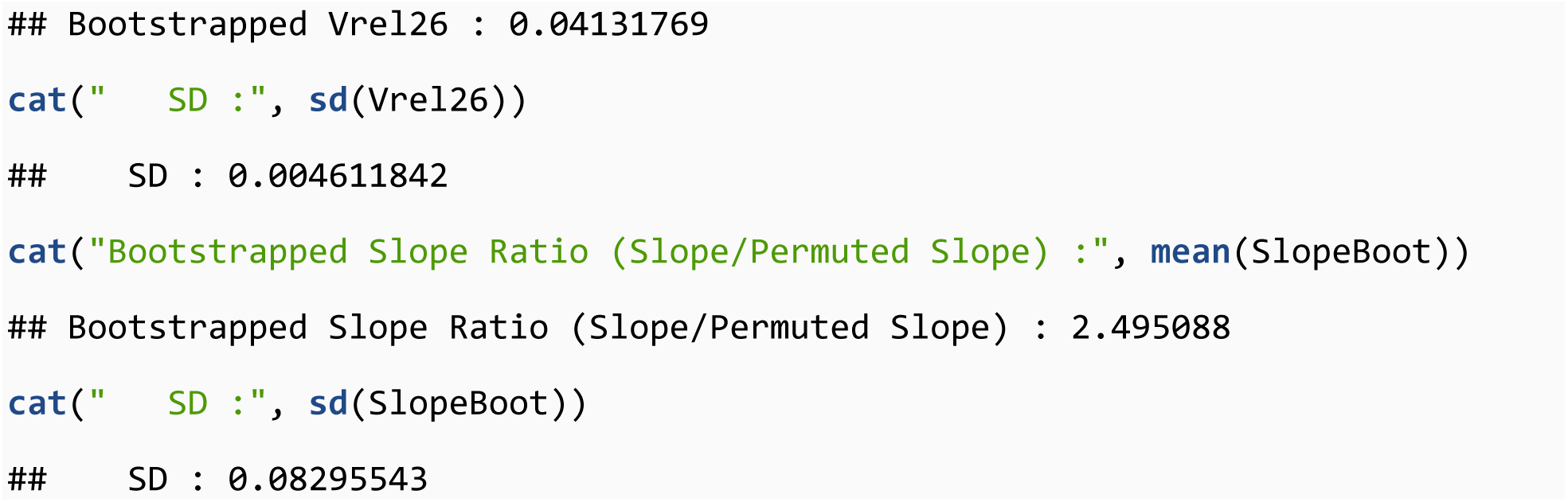

